# An atlas of the human metabolome

**DOI:** 10.64898/2026.05.21.726638

**Authors:** Jeremy K. Chan, Nicholas S. Ly, Olivia Taverniti, William D. Gwynne, Brandon Y. Lieng, Vanessa Affe, Verne T. Urquhart-Cox, Sophia M. Alonzi, Mathula Muhundan, Alexandra J. Denhart, Landon J. Edgar, Andrew T. Quaile, J. Rafael Montenegro-Burke

## Abstract

Despite the emergence of cellular atlases like the Human Protein Atlas, no equivalent atlas exists for the human metabolome. Here, we present the Human Metabolome Atlas (HMA, <u>hma.ccbr.utoronto.ca</u>), a comprehensive map containing metabolomic profiles of 70 human cell lines across 22 tissues. With an ∼8-fold increase in coverage compared to other resources, the HMA contains quantitative data for 1768 metabolites at the highest identification confidence, encompassing over 50 lipid classes and a broad range of metabolic pathways. This constitutes the most extensive human metabolomic atlas available. Leveraging the HMA, we identified specific metabolic regulation within pathways and cell types and characterized metabolic processes like glycosylation and ferroptosis. Lastly, we developed a publicly available, interactive web-portal to facilitate custom data analysis for the broader scientific community.

## INTRODUCTION

Metabolomics offers a comprehensive view of the small molecules that participate in life-sustaining chemical reactions (*1*). These metabolic reactions often vary by cell type and fluctuate in response to intra- and extra-cellular conditions while concomitantly exerting their own independent effects. Dysregulation of these evolutionarily balanced processes is fundamental to the outcomes of many diseases such as cancer (*2*, *3*). Consequently, metabolomics continues to grow in popularity as a tool for investigating the underlying mechanisms of disease (*4*, *5*). The chemical complexity of the metabolome, however, particularly in comparison to genetic sequence based ‘-omics’, has limited the rate at which its depth and heterogeneity can be characterized (*6*, *7*). As such, our knowledge of the human metabolome remains relatively rudimentary, although continued technological advancements have made obtaining this data at a meaningful scale increasingly feasible (*8*).

Resources like the Human Protein Atlas (HPA) and the Cancer Dependency Map (DepMap) illustrate how large-scale compendia of human biology across cells and tissues can be used as powerful tools to understand cellular function and processes (*9–11*). Large scale human metabolomics resources (e.g., HMDB, METLIN, MoNA) currently available serve as invaluable encyclopedic repositories of metabolites and their physiochemical properties; but offer limited capacity for quantitative comparative analysis between different cells and tissues (*12–14*). Among the resources with quantitative information, the data available are often limited to non-human specimens (e.g., mouse) or to specific human-derived tissues or samples (e.g., heart, plasma) (*15–17*). Furthermore, while the closest metabolome resource analogous to the HPA provides extensive biological coverage, its ability to deeply interrogate metabolic processes is restricted by its limited metabolome coverage (*18*). Hence, there remains a clear need for a resource that captures the full complexity of the human metabolome across a wide range of metabolites and cell types.

Here, we present the Human Metabolome Atlas (HMA)—a comprehensive dataset that contains the metabolomic and lipidomic profiles of 70 human cell lines across 22 tissue types. We employed complementary state-of-the-art liquid chromatography mass spectrometry (LC-MS) approaches to achieve exceptional metabolome coverage and relative quantitation of 1768 metabolites and lipids at the highest identification confidence possible. This resource represents the largest and broadest human metabolome compendium and provides an ∼8-fold increase in metabolome coverage compared to its closest available peer. Leveraging this expansive resource, we uncovered tissue-specific metabolomic signatures and global associations among metabolites and lipids. Our analyses also revealed tissue-specific enrichment of nucleotide intermediates and triglycerides in hematological cell lines, which have functional implications for cellular processes like glycosylation and ferroptosis. Finally, we built a publicly available, interactive web-portal (<u>hma.ccbr.utoronto.ca</u>) to facilitate exploration of the HMA dataset. The web-portal is a unique platform especially designed for metabolomics and lipidomics data, which offers advanced and customizable data analysis capabilities like hierarchical clustering and Fatty Acid Composition Heatmaps (FACHs) (19). Together, the HMA represents a powerful resource and discovery platform that harnesses metabolomics to deepen our understanding of human biology.

## RESULTS

### Development of the Human Metabolome Atlas (HMA)

To ensure high confidence and broad coverage in the HMA, we developed a reproducible experimental pipeline for the large-scale acquisition of metabolite (polar small molecules) and lipid (non-polar small molecules) profiles. **(Fig. 1A and fig. S1)**. The development of the HMA consisted of cell culture, metabolite and lipid extraction, LC-MS analysis, and data analysis. Briefly, 70 human cell lines from 22 tissue types were cultured in replicate (n = 5) for analysis by LC-MS **(Fig. 1B and table S1).** Each cell line was also cultured for bulk RNA-sequencing to supplement the metabolomics and lipidomics data **(table S2)**. For sample preparation, we used a biphasic methyl-tert-butyl ether (MTBE) method that enables the simultaneous extraction of metabolites and lipids (*15*, *20*, *21*) **(fig. S1)**. Moreover, to ensure reproducibility and quality control throughout the experimental workflow, we implemented multiple measures for assessing and minimizing variation. These included cell culture batch controls, isotopically labeled standards, pooled LC-MS quality controls, and sample randomization spanning the experimental workflow, from extraction to injection **(fig. S1, see Methods)**.

**Fig. 1.**
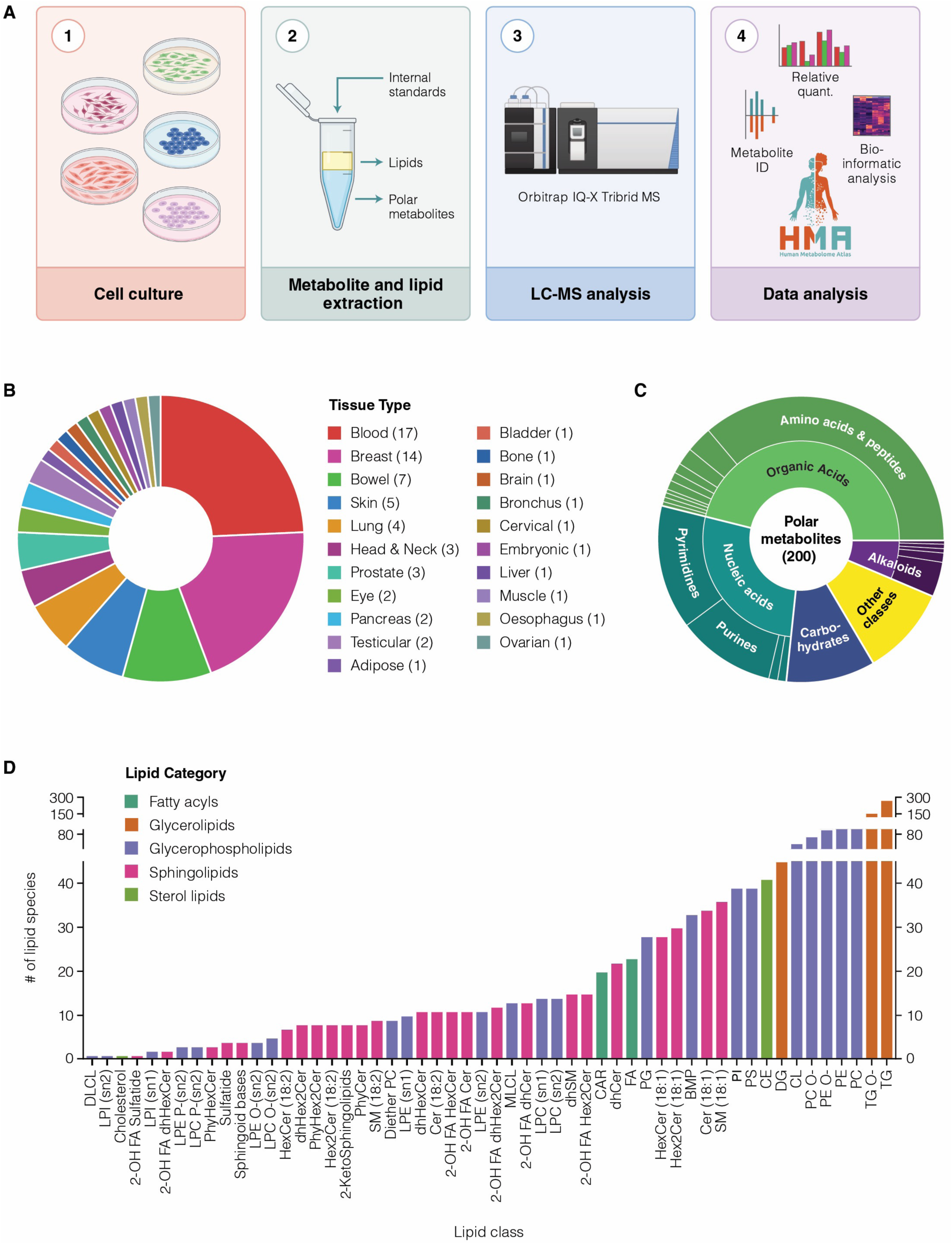
Development of the Human Metabolome Atlas (HMA). **(A)** Experimental workflow for the HMA. For further details, see Fig. S1. **(B)** Tissue distribution of the 70 human cell lines profiled for the HMA. **(C)** Polar metabolites profiled in the HMA grouped by metabolite classes. **(D)** Lipid species profiled in the HMA grouped by lipid class and colored by lipid category.

To maximize metabolome and lipidome coverage, cell lines were analyzed using three complimentary LC-MS methods. Metabolite extracts were analyzed using two different hydrophilic interaction liquid chromatography (HILIC) MS methods while lipid extracts were analyzed using a reversed-phase liquid chromatography MS method. Metabolites and lipids were identified at high confidence and according to the guidelines established by the Metabolomics Standards Initiative (MSI) (*22*, *23*). Metabolites were identified using experimental data that matched authentic reference standards (MSI level 1) **(fig. S2A)**. For lipids, most identifications were reported in sum composition notation, accurately reflecting the level of structural resolution achieved. Lipids were identified at a minimum confidence level of MSI level 2 using a combination of accurate mass and MS/MS fragmentation. To further validate lipid identities, retention times (RTs) derived from commercially available or in-house, synthesized standards were also matched and extrapolated to measured lipid RTs **(fig. S2B)** (*24*, *25*). Collectively, for the HMA, we identified and quantified the relative levels of 1768 molecules, consisting of 200 metabolites and 1568 lipids **(table S3-4)**. The 200 metabolites encompass many well-characterized metabolic pathways including amino acid and nucleotide metabolism **(Fig. 1C)**. Likewise, with 1568 lipid species across 54 lipid classes, the HMA contains data for a wide range of lipids including glycerophospho-, glycerol-, and sphingolipids **(Fig. 1D)**. Given the broad chemical coverage and level of confidence in metabolite identifications, the HMA constitutes the most comprehensive and rigorous human metabolomics atlas currently available.

A major aim of the HMA was to develop a metabolomics-based resource that was relevant and practical for the scientific community. Thus, we chose to culture our cell lines in commonly used or vendor-recommended cell culture media. This approach introduces a potential confounding variable where differences in media composition could substantially alter the metabolome (*26*). To address this, we assessed the effects of various common basal media conditions on the metabolome. We cultured three cell lines (K562, HeLa, and BJ) in either their recommended basal cell culture medium (IMDM, DMEM, and EMEM, respectively) or RPMI-1640 (RPMI) and profiled their metabolomes and lipidomes. Hierarchical clustering of combined metabolomic and lipidomic profiles showed that RPMI-cultured cell lines cluster according to their original cell line identity and not by growth media condition **(fig. S3A)**. This trend also held true when the dataset was restricted to only metabolites, which are the only variable components in the different basal media **(fig. S3B)**. These observations suggest that although differences in media composition should be considered as an important experimental variable, the overall metabolic characteristics of each cell line in the HMA are maintained. Importantly, this allows for comparisons of the metabolomes across all 70 cell lines in our atlas, while maintaining in optimal growth conditions for each line.

### Global analysis of cell lines reveals context-specific metabolomic and lipidomic signatures

To characterize the global metabolic landscape and to gain insight into the metabolic regulation of cell lines within the HMA, we first examined tissue-level correlations of metabolomic and lipidomic profiles. Hierarchical clustering of Spearman’s correlation coefficients revealed that developmentally-related tissues shared similar metabolomic and lipidomic profiles. For instance, cell lines from endoderm-derived tissues of the digestive system (bowel, pancreas, and oesophagus) were strongly correlated with each other **(Fig. 2A)** (*27*). Similar correlations were also observed among mesoderm-derived gonadal tissues (testicular and ovarian) (*28*).

**Fig. 2.**
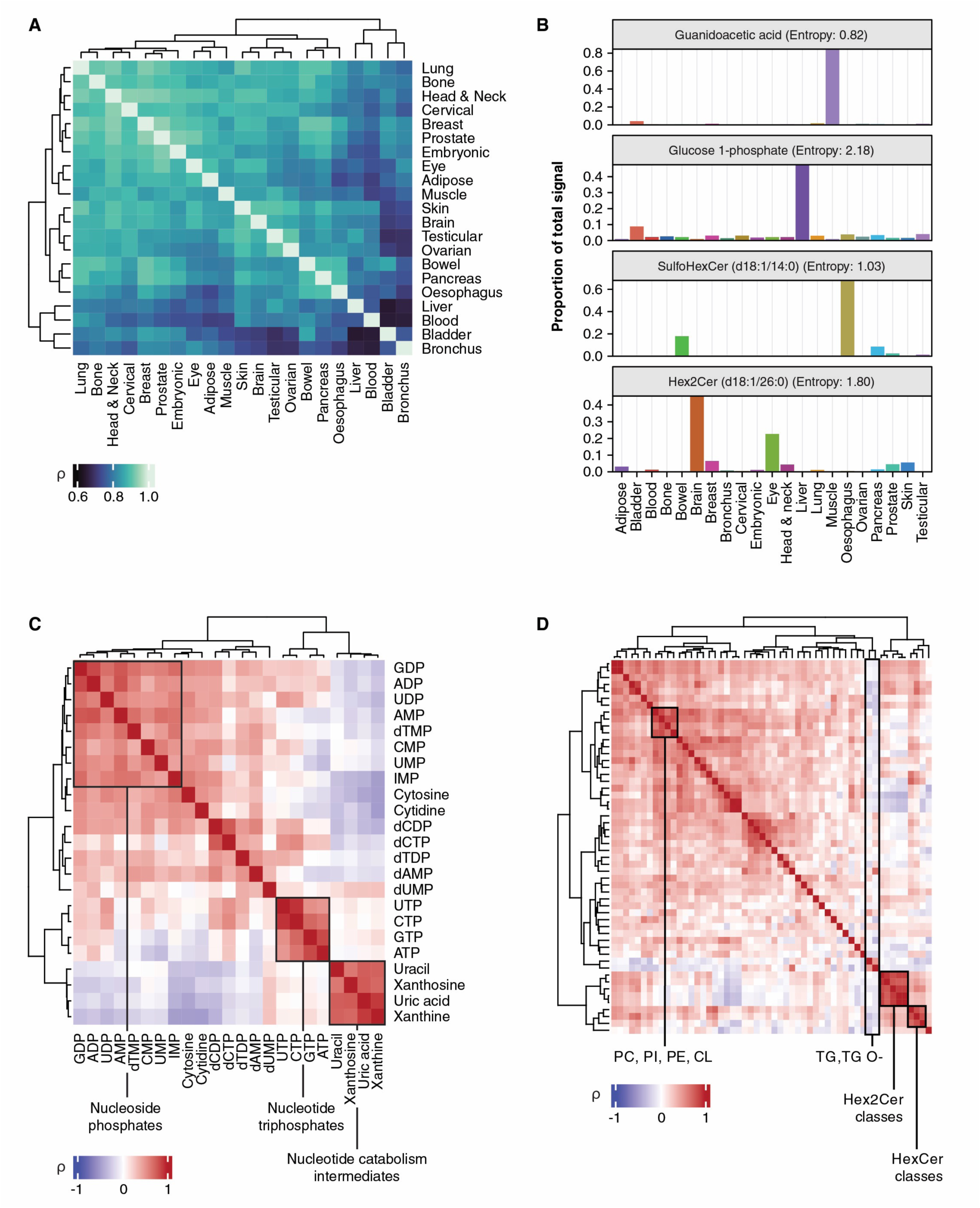
Global analysis of metabolomic profiles in the HMA. **(A)** Correlation heatmap of metabolomic profiles grouped by tissue of origin. Data are presented as Spearman’s rho (ρ) of 1768 metabolites and lipids across 21 tissue types. **(B)** Examples of tissue-enriched metabolites and lipids identified using Shannon entropy. Data are presented as the proportion of total signal for a particular metabolite within a given tissue. **(C)** Correlation heatmap of nucleotide pathway metabolites. Data are presented as Spearman’s rho (ρ) of mean metabolite abundance across 70 cell lines. **(D)** Correlation heatmap of the summed abundance of individual lipid classes across 70 HMA cell lines. Data is presented as Spearman’s rho of lipid classes across 70 cell lines (ρ). Boxes depict specific groups of related lipid classes. Abbreviations: phosphatidylcholine, PC; phosphatidylethanolamine, PE; cardiolipin, CL; phosphatidylinositol, PI; triglyceride, TG; ether-linked triglyceride, TG O-; hexosylceramide, HexCer; dihexosylceramide, Hex2Cer.

This connectivity between tissue types prompted us to identify metabolites and lipids that not only underlie these correlations but could also serve as tissue-specific metabolic markers. To this end, we used Shannon entropy (SE), which has previously been used to quantify feature specificity in metabolomics and other biological datasets (*29*, *30*). Low SE indicates high feature specificity within a specific group (i.e., enrichment) while high SE suggests that the dataset is more evenly distributed across many groups. Using this approach, SE was calculated for each metabolite on a tissue basis and those with values in the bottom quartile were defined as “tissue-specific” **(fig. S4A and table S5)**. Closer examination of low SE metabolites and lipids led to the identification of several tissue-specific metabolic signatures **(Fig. 2B)**. For instance, we observed enrichment of guanidoacetic acid, a precursor of creatine, in the muscle-derived cell line RH-30. This aligns with our understanding of muscle function and muscle cells, which heavily rely on creatine for ATP cycling (*31*, *32*). Similarly, glucose-1-phosphate (G1P), an intermediate of glycogenesis/glycogenolysis, was found to be enriched in the liver-derived HepG2 cell line. This enrichment is consistent with the fact that the liver is the primary site for glycogenesis/glycogenesis (*33*). Likewise, tissue-specific enrichment was observed across multiple glycosphingolipid classes, including sulfatides and dihexosylceramides (Hex2Cers). Consistent with previous findings, sulfatides were enriched in digestive system-related cell lines (e.g., bowel, oesophagus, pancreas) **(fig. S4B-D)** (*34*). Likewise, cerebrosides like Hex2Cers were enriched in the glioblastoma cell line U138-MG **(fig. S4E-G).** This is consistent with previous studies that have emphasized the importance of cerebrosides for brain function (*35*, *36*). Together, these data highlight the utility of the HMA in identifying metabolites that drive tissue-tissue relationships and serve as tissue-specific markers for different metabolic processes.

Although these correlations provided significant insight on tissue-tissue relationships, it also prompted us to ask if these metabolomic and lipidomic profiles could be distinguished based on other biological or molecular classifiers. To answer this, we performed principal component analysis (PCA) on the combined metabolomic and lipidomic profiles of all HMA cell lines. The PCA highlighted two groups of cell lines across PC1 that broadly clustered based on hematological (myeloid and lymphoid) or non-hematological tissues of origin **(fig. S5A)**, in line with previous reports (*18*). Further examination into specific metabolites and lipids that drive these clusters revealed elevated levels of nucleotide intermediates **(fig. S5B)** and triglycerides (TGs) **(fig. S5C)**. Concomitant trends between metabolites or lipids within the same metabolic pathway (e.g., nucleotide metabolism, TG species) suggest their levels are coregulated or interdependent. To identify coregulated metabolites, we first examined the Spearman’s correlations between pairs of metabolites across the HMA. Using this approach, we identified distinct correlations between nucleotide metabolism intermediates. Specifically, we observed strong correlations between nucleoside mono- (NMPs) and diphosphates (NDPs) (e.g., AMP, UDP, etc.) **(Fig. 2C)**. Similarly, nucleoside triphosphates (NTPs) (e.g., UTP) were strongly correlated with one another; however, NTP levels did not correlate to their respective NMPs and NDPs. In contrast, metabolites related to nucleotide catabolism (e.g., uracil, xanthosine) were strongly correlated with one another but negatively correlated with NMPs and NDPs. Moreover, the same analysis was performed for the lipidome at the lipid class level to gain an understanding of coordinated trends across structurally and functionally distinct lipids. Pairwise lipid class correlations revealed strong correlations between structurally similar lipid classes like phosphatidylcholines (PCs), phosphatidylethanolamine (PEs), and phosphatidylinositols (PIs), which are all prominent components of cellular membranes (*37*). Similarly, sphingolipid classes like hexosylceramides (HexCer) and Hex2Cers were strongly correlated with one another **(Fig. 2D and fig. S6)**. In contrast, we found that TGs and ether-linked TGs (TG O-) were strongly correlated with one another; however, neither correlated strongly with any of the other 52 lipid classes. Taken together, these findings demonstrate how the broad biological and metabolic coverage of the HMA can be leveraged to gain deeper insights into the highly orchestrated nature of metabolic processes in different metabolic pathways.

### Inferring glycosylation phenotypes from HMA metabolomic profiles

To better understand the metabolic differences that distinguish hematological and non-hematological cell lines, we examined the most significantly altered metabolites and their pathways **(fig. S5A)**. Comparison between these two groups revealed that nucleotide intermediates were amongst the most differentially abundant metabolites **(fig. S5B and table S6)**. A clustered heatmap of these pathway intermediates revealed a distinct elevation in NMPs and NDPs in hematological cell lines compared to non-hematological cell lines **(Fig. 3A)**. In contrast, metabolites related to nucleotide catabolism (e.g., uracil) were detected at significantly lower levels in hematological cell lines. This stark contrast between closely related metabolites indicates that nucleotide pools are tightly regulated and may be amassed for specific cellular processes in hematological cell lines. To determine how these nucleotide pools are being regulated, we turned to gene expression data derived from the bulk RNA-sequencing of HMA cell lines. By correlating metabolite ratios with mRNA transcript levels, we identified substrate-product relationships that potentially explain the underlying variation in nucleotide metabolism. We found that the uracil to uridine ratio was positively correlated with the expression of uridine phosphorylase 1 (*UPP1*), an enzyme that facilitates the conversion of uridine to uracil **(Fig. 3B)**. Compared to non-hematological cell lines, hematological cell lines tended to have lower ratios of uracil to uridine along with lower expression of *UPP1*, which may explain the accumulation of nucleotides. When we expanded this analysis to genes and metabolites involved in *de novo* pyrimidine synthesis, we also observed no significant correlation between upstream pyrimidine metabolites and corresponding genes **(fig. S6A-D)**. Taken together, these results suggest that the accumulation of these pyrimidine intermediates may reflect lower levels of catabolism, implying that these metabolites could be accumulating for other cellular processes.

**Fig. 3.**
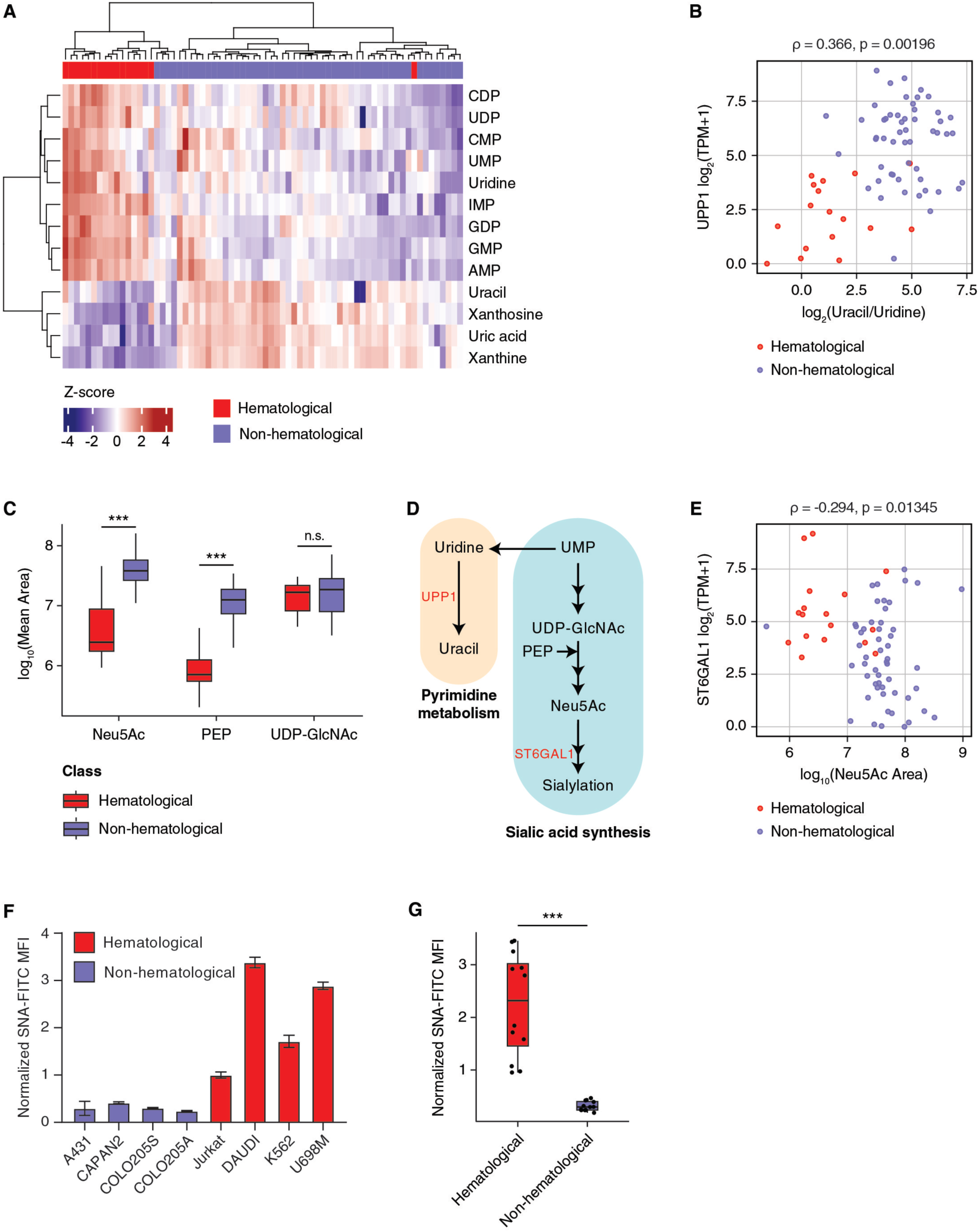
Elevation of nucleotide intermediates in hematological cell lines is associated with increased cell surface sialylation. **(A)** Clustered heatmap of nucleotide intermediates in all HMA cell lines. Red bars represent hematological cell lines while purple bars represent non-hematological cell lines. Rows represent metabolites related to nucleotide metabolism. Data are presented as Z-score of log10-transformed peak areas of specified metabolites. All metabolites shown are statistically significant between hematological and non-hematological cell lines (adj. p-value <0.05, Mann-Whitney U test, Benjamini-Hochberg correction). **(B)** Scatter plot of the log2 ratio of uracil to uridine abundances compared to UPP1 expression of cell lines in the HMA. Individual points represent HMA cell lines colored based on tissue of origin. Correlation was calculated using Spearman’s correlation. **(C)** Boxplots depicting the mean abundance of sialylation precursors in hematological cell lines and non-hematological cell lines. Data are presented as log10 mean peak area. *** denotes significance p<0.0001 (Mann-Whitney U test, Benjamini-Hochberg correction). Abbreviations: Uridine diphosphate-N-Acetylglucosamine, UDP-GlcNAc; phosphoenolpyruvic acid, PEP; N-Acetylneuraminic acid, Neu5Ac. **(D)** Summarized metabolic pathway of pyrimidine metabolism linked to sialic acid synthesis. **(E)** Scatter plot of log10 abundance of Neu5Ac compared to ST6GAL1. Correlation was calculated using Spearman’s correlation. **(F)** Quantification of α-2,6 sialylation across various cell lines in the HMA. Data are presented as normalized SNA lectin MFI values with signal normalized to the mean normalized lectin MFI of the Jurkat cell line (n= 3). **(H)** Statistical analysis of cell surface α-2,6 sialyation grouped tissue of origin. Data are presented as mean normalized MFI values from panel (F). *** denotes statistical significance by Mann-Whitey U Test (p-value <0.0001).

Functionally, metabolites related to nucleotide metabolism are commonly associated with essential cellular processes like DNA and RNA synthesis. In addition, nucleotide intermediates are also involved in cell signaling and glycosylation, which are extremely heterogeneous cellular processes that vary widely in terms of regulation and activity based on cell type (*38*, *39*). Glycosylation is of particular interest as we previously showed that the depletion of pyrimidines is a metabolic vulnerability of group 3 medulloblastoma cells and can result in lower levels of protein glycosylation (*40*). Based on this, we surmised that the elevation of nucleotide intermediates may be reflected in glycosylation and its metabolic precursors. Hence, to understand the relationship between pyrimidine metabolism and glycosylation, we compared the abundance of glycosylation precursors between hematological and non-hematological cell lines in the HMA. Interestingly, we found that levels of *N*-acetylneuraminic acid (Neu5Ac) and phosphoenolpyruvic acid (PEP) were significantly lower in hematological cell lines **(Fig. 3C)**. Notably, Neu5Ac and PEP serve as precursors for sialylation, a type of glycosylation commonly found on the surface of human cells **(Fig. 3D)** (*41*, *42*). In contrast, the levels of uridine diphosphate *N*-acetylglucosamine (UDP-GlcNAc)—a substrate for other glycosylation pathways and an upstream precursor of Neu5Ac—were comparable between hematological and non-hematological cell lines. Subsequent investigation into the genes involved in sialylation revealed that Neu5Ac was negatively correlated to the expression of *ST6GAL1*, a sialyltransferase **(Fig. 3E)**. As Neu5Ac is a precursor for sialylation, we hypothesized that *ST6GAL1* consumes Neu5Ac at higher levels for sialylation in hematological cell lines. Accordingly, this should be reflected in levels of cell surface sialylation. To validate this, we used flow cytometry to quantify cell surface sialylation in multiple HMA cell lines using a fluorescent *Sambucus nigra* (SNA) lectin that binds α2-6 linkages between sialic acids (e.g., Neu5Ac) and galactose, generally in the context of N-linked glycans **(Fig. 3F and fig. S6E)** (*43*). Indeed, hematological cell lines exhibited significantly higher levels of α2-6 sialylation than non-hematological cell lines, consistent with higher *ST6GAL1* activity and increased consumption of sialylation precursors **(Fig. 3G)**. Together with the accumulation of nucleotide intermediates, this also suggests that the elevation of nucleotide intermediates is necessary to replenish precursors like Neu5Ac and PEP. Ultimately, these results demonstrate how the HMA can be used beyond investigating metabolism and can be extended to other cellular processes like protein sialylation.

### Identifying ferroptosis-susceptible cell lines from HMA lipidome profiles

The striking metabolomic differences between hematological and non-hematological cell lines prompted us to further explore differences in their lipidomes. As previously mentioned, we observed significant differences in TG species between hematological and non-hematological cell lines **(fig. S5C)**. Closer examination revealed that many of these elevated TG species were highly unsaturated (>3 double bonds) **(fig. S7A)**. To better understand these highly unsaturated TGs, we used FACHs (*21*). FACHs deconvolute lipid sum compositions into acyl chain length and double bond content, so the proportional abundance of hundreds of individual lipid species can be easily visualized at the lipid class level. As an analytical tool, FACHs provide insight on how individual fatty acids are distributed across a lipid class. Comparison between hematological and non-hematological cell lines showed that hematological cell lines contained longer and more unsaturated TGs **(fig. S7B-D)**. This was also evident when visualizing TG acyl chain length and double bond content at the individual cell line level **(Fig. 4A and table S8)**. Overall, these findings suggest that TG acyl chain length and acyl double bond profiles are lipidomic signatures that can be used to distinguish hematological from non-hematological cell lines.

**Fig. 4.**
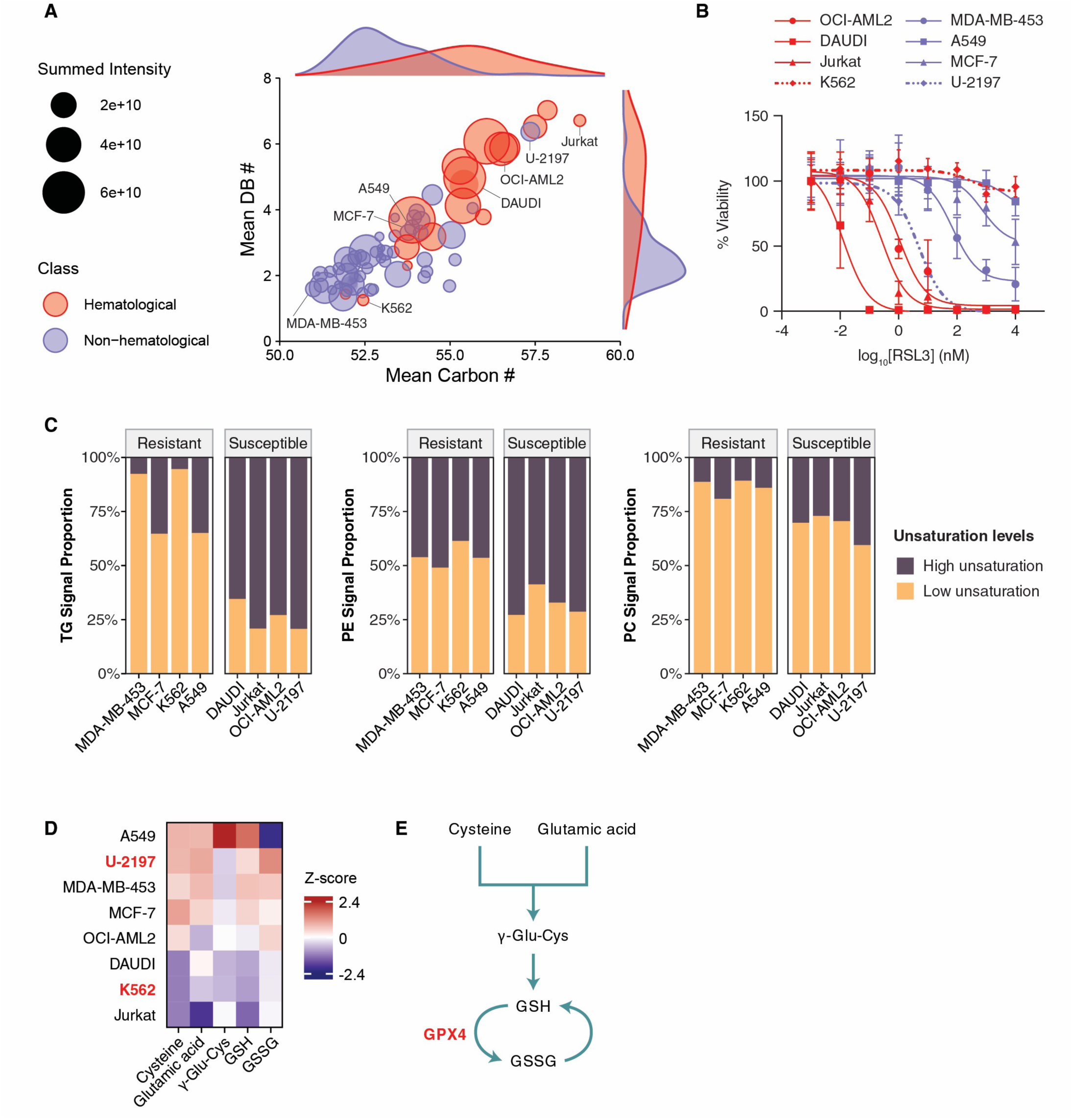
Identifying ferroptosis-susceptible cell lines from HMA lipidome profiles. **(A)** Joint plot representing the distribution of triglycerides (TGs) across the 70 HMA cell lines. Scatter plot shows the mean mean number of carbons and mean number of double bonds in the acyl chains of TGs for each cell line. Size of each point is scaled to the summed intensity of all TG species in each cell line. Marginal plots depict the distribution of weighted carbon number and weighted double bond number values. **(B)** Dose-response curve of cell lines treated with RSL3 for 24 hrs. Signal was measured using PrestoBlue. Adherent cell lines are colored in blue while suspension cell lines are colored in orange. Data are shown as cell viability normalized to vehicle-control (mean ± s.d., n =4-5) **(C)** Stacked bar plot depicting the proportional abundance of lipid species with high unsaturation compared to those with low unsaturation for RSL3-tested cell lines. Each plot depicts a specific lipid class. High unsaturation lipid species are defined as lipid species that have double bond numbers greater than the number of acyl chains in a given lipid class. Low unsaturation lipid species are defined as lipid species that have a double bond number less than or equal to the number of acyl chains in a particular lipid class. **(D)** Heatmap of metabolites involved in glutathione metabolism for RSL3-tested cell lines. Data are presented as scaled Z-score abundance for each metabolite. **(E)** Schematic of glutathione metabolism pathway.

The pronounced difference in acyl chain double bond composition is particularly noteworthy as the addition or removal of a single double bond can have distinct biological implications (*44*). This is apparent when examining the effects of saturated fatty acids (SFAs), monounsaturated fatty acids (MUFAs), and polyunsaturated fatty acids (PUFAs), which have been reported to have distinct effects in inflammation and cancer (*45–47*). Notably, hematological cell lines, on average, have TGs with ∼2 more double bonds than non-hematological cell lines, which potentially represents a shift from SFA- to PUFA-containing species **(fig. S7D)**. In comparison, the difference in double bond content for other lipid classes was lower in magnitude, with both PCs and PEs within hematological cell lines having on average only ∼0.75 more double bonds than non-hematological cell lines **(fig. S7E-F)**. Given this, we aimed to investigate the functional implications associated with these variations in double bond profiles. In particular, we identified ferroptosis susceptibility as a phenotype of interest as it has been heavily associated with PUFA-containing lipids (*48*). As a cellular process, ferroptosis is characterized by the accumulation of lipid peroxides derived from PUFA-containing lipids such as PUFA-phospholipids (PUFA-PLs), leading to a form of programmed cell death (*48*, *49*). Cells with high levels of these PUFA-PLs have also been found to be more susceptible to ferroptosis (*50*, *51*). More recently, there has been growing evidence that other lipid classes like PUFA-TGs also contribute to ferroptosis (*52*, *53*). Based on this, we hypothesized that we could identify ferroptosis-susceptible cell lines based on TG profiles. To test this, we selected cell lines with a range of TG double bond profiles and performed a dose-response assay using a potent glutathione peroxidase 4 (GPX4) inhibitor and ferroptosis inducer (1S,3R)-RSL3 (RSL3) (*54*). Indeed, we found that RSL3-susceptible cell lines were enriched for PUFA-TGs relative to RSL3-resistant cell lines. This effect was independent of tissue of origin (hematological vs non-hematological), as indicated by the K562 and U-2197 cell lines (**Fig. 4B and fig. S7G)**. K562 is a hematological cell line but has a similar TG profile to an average non-hematological cell line with its high proportion of saturated TGs, which may explain its resistance to RSL3 **(Fig. 4A and fig. S8A)**. Inversely, U-2197 is a non-hematological cell line but has a similar TG profile to an average hematological cell line with its high proportion of unsaturated TGs, consistent with RSL3 susceptibility **(Fig. 4A and fig. S8B)**. Taken together, these data further support the hypothesis that unsaturated TGs play a significant role in determining ferroptosis susceptibility. Beyond TGs, we also examined whether other lipid classes could be used to determine ferroptosis susceptibility. Closer investigation revealed significantly higher proportions of PUFA-PCs and PUFA-PEs in RSL3-susceptible cell lines **(Fig. 4C and fig. S9)**. This is consistent with previously work that highlighted the role of unsaturated phospholipids (PLs) in ferroptosis (*52*, *55*). Notably, there was a greater difference in the proportional abundance of PUFAs in TGs (3-fold) than in PLs (2-fold), suggesting TGs may be a better predictor of ferroptosis susceptibility **(Fig. 4C)**.

While PUFA-lipids are associated with ferroptosis susceptibility, other metabolic pathways and metabolites also confer ferroptosis resistance. One of the main mechanisms of ferroptosis resistance is facilitated GPX4 and the glutathione metabolic pathway (*54*). GPX4 protects cells from ferroptosis by reducing lipid peroxides to lipid alcohols through the consumption of reduced glutathione (GSH) (*54*, *56*). Interestingly, when we examined the levels of metabolites like GSH, we found them elevated in non-hematological cell lines compared to hematological cell lines, independent of TG profiles **(Fig. 4D-E)**. This suggests that glutathione pathway intermediates contribute to ferroptosis resistance, but do not alone predict ferroptosis susceptibility. This is illustrated by the K562 cell line, which has lower levels of glutathione pathway intermediates but are resistant to RSL3-induced ferroptosis. Collectively, these data identified unsaturation levels in TGs as a highly specific maker for ferroptosis susceptibility and demonstrates the functional utility of the broad metabolic coverage of the HMA.

### The Human Metabolome Atlas Discovery Portal

One of the primary objectives of the HMA was to provide an accessible, scalable, and functional resource for the broader scientific community that could be used for metabolomics-based biological discovery. To do this, we developed a publicly accessible web portal (<u>hma.ccbr.utoronto.ca</u>) with the data for all 1768 metabolites and lipids across all 70 cell lines profiled. This specialized web-portal provides detailed information on each metabolite (e.g., molecular structure, molecular formula, monoisotopic mass, *m/z*, etc.) as well as links to other invaluable community resources such as HMDB, SwissLipids, KEGG, and PubChem **(Fig. 5A)** (*12*, *57*, *58*). We have also extensively curated cell line metadata including tissue of origin, disease type, treatment information, mutations, ancestry, and sex. In addition to this information, the defining advantage of the HMA is its unique ability to facilitate user-driven exploration of the human metabolome. The is done through customizable data visualization tools for comparing metabolites and lipids across cell lines and tissue types **(Fig. 5B)**. For more complex data analysis, we provide tools to perform hierarchical clustering, dimensionality reduction, and two-class comparisons (e.g., volcano plots). To demonstrate the utility of these tools, we show a proof-of-concept comparison between luminal and basal-like breast cancer cell lines that reveals significantly higher levels of hydroxylated ceramides in luminal breast cancer cell lines **(Fig. 5B**). Another example of this is the use of boxplots on the web portal to highlight elevated levels of G1P in the liver **(Fig. 5C)**. Amongst the data visualization tools available, FACHs are one of the most powerful as they capitalize on the extensive coverage of the HMA to summarize the distribution of acyl chain carbons and double bonds across lipid classes and cell lines in a 2-dimensional plot **(Fig. 5D)** (*21*). These give an intuitive overview of the varied distributions and expected complexity of lipids for the HMA cell lines, and provide context needed to understand the significance and magnitude of changes that may be observed in lipidomics datasets. Taken together, the built-in tools of the HMA combine deep metabolome and lipidome coverage, ease of use, and flexible data analysis options to provide a robust metabolomics-based discovery platform for the broader scientific community.

**Fig 5.**
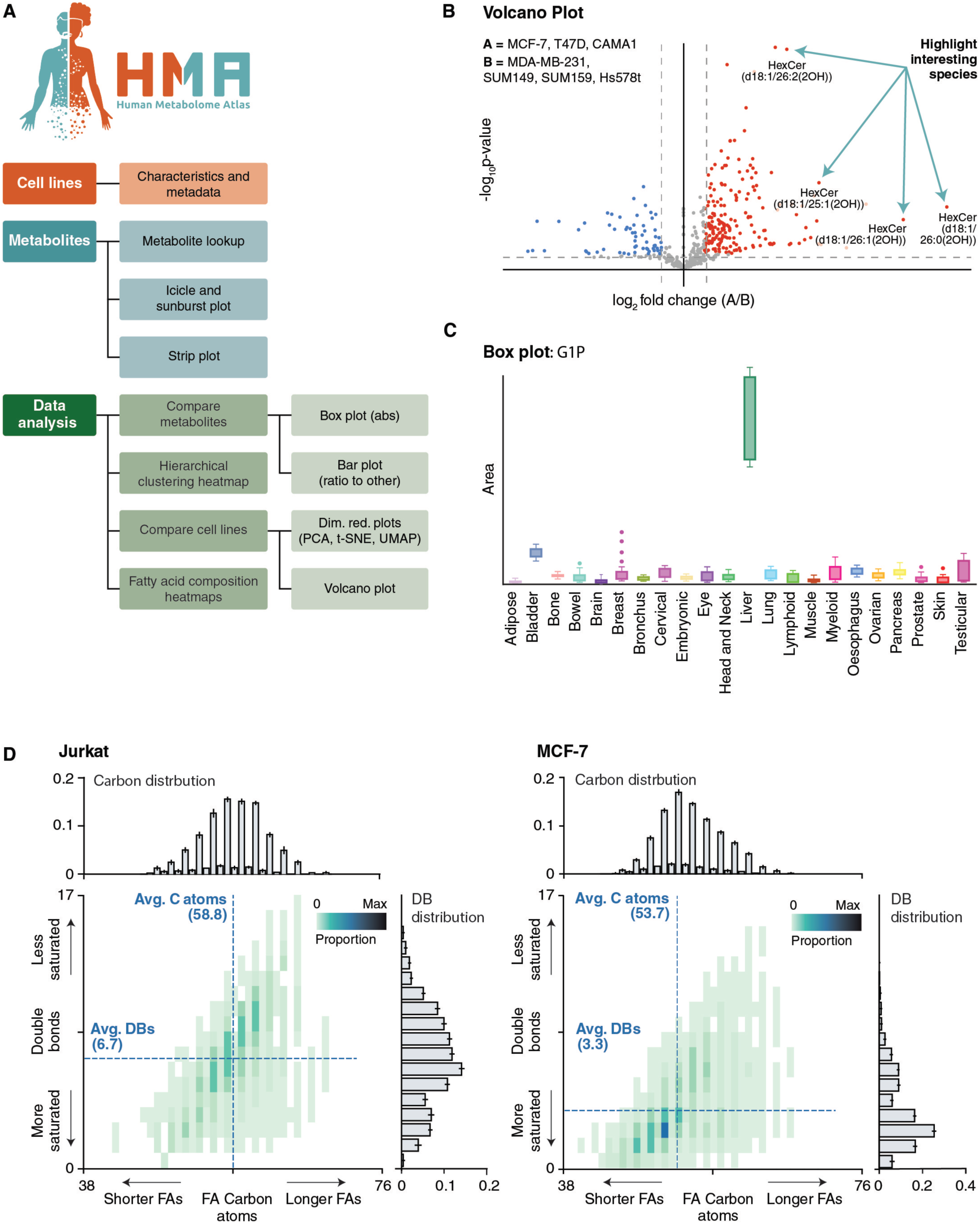
The Human Metabolome Atlas discovery portal. **(A)** Site-map of the v1.0 release of the HMA web-portal. The site is navigated via three primary areas: “Cell Lines” – for browsing of information about the cell lines in the HMA, such as tissue origin, sex, ethnicity, disease etc.; “Metabolites” – for displaying data pertaining to one or more metabolites, such as chemical formulas and measured abundance across cell lines/tissues; and “Data Analysis” - for performing more complex data analysis tasks such as statistical comparisons of groups of cell lines, hierarchical clustering and principal component analyses. **(B)** Example volcano plots showing log2 (A/B) fold-changes and statistical significance for metabolites between user selected breast cancer cell lines. **(C)** Example box plot comparing measured peak areas for G1P across all cell lines, grouped by tissue type. **(D)** Example TG FACHs that compare the proportional distribution of acyl carbon atoms and double bonds between two cell lines.

## DISCUSSION

The recent proliferation of metabolomics in biological research has highlighted the need for ‘atlas-like’ resources for expediting scientific discovery (*59*). To address this gap, we present the HMA (<u>hma.ccbr.utoronto.ca</u>)—an interactive and publicly accessible resource containing the relative levels of 1768 metabolites and lipids across 70 human cell lines. The HMA stands out among other metabolomics resources due to its extensive chemical coverage across a diverse collection of cell types. Notably, the HMA contains data for ∼8 times more metabolites and lipids than its closest comparable at the highest identification confidence level possible (*18*).

Using this resource, we reaffirm tissue-specific metabolite signatures while also identifying novel relationships between molecules. Further analysis into the HMA dataset demonstrated that the metabolomic and lipidomic profiles of human cell lines can be broadly classified based on tissue of origin (hematological vs non-hematological). We demonstrated that hematological cell lines regulate nucleotide metabolism to support higher levels of cell surface sialylation. At the lipidome level, our work further demonstrates the role of PUFA-TGs in ferroptosis and highlights how TG profiles from the HMA can be used as a signature to predict cell line susceptibility to RSL3-induced ferroptosis (*53*).

There are some limitations in the HMA that should be noted. First, the current iteration of the HMA contains data for 70 human cell lines. From the thousands of cell lines available for biomedical research, we profiled a representative subset that would robustly capture the diversity of human tissue types. Second, the HMA provides a snapshot of the metabolome and lipidome and does not capture the dynamics of metabolic processes (e.g. flux analysis). Lastly, the chemical complexity and concentration range of the human metabolome and lipidome precludes the comprehensive detection of all metabolites and lipids, as no single analytical approach can capture its entirety at high confidence. This is best illustrated through the lipids in the HMA, which are mostly represented in sum composition as we cannot consistently deconvolute acyl chain composition or double bond positions for datasets at such a large scale. Thus, potential extensions include the development of methodologies that enable the reliable and high throughput detection of lipids at higher structural resolution. Collectively, the HMA uniquely combines extensive metabolome and lipidome coverage with specialized interactive data exploration to expedite our understanding of the human metabolome and drive biological discovery.

## MATERIALS AND METHODS

### Cell lines and culture conditions

#### Procurement of cell lines

Commercially available cell lines of malignant and normal tissue origin were received as in-kind gifts from collaborating labs or procured from the American Type Culture Collection (ATCC), Evercyte and The Leibniz Institute DSMZ **(table S10)**. Cells were expanded according to the media conditions and seeding densities recommended by each supplier or collaborating lab. PDC005 is a patient-derived head and neck squamous cell carcinoma cell line provided in-kind by Prof. Daniel Schramek’s lab.

#### Cell line quality control

Cell lines were determined to be free of mycoplasma infection *via* Lonza MycoAlert® Detection Kit (Lonza Biosciences, #LT07-318). In brief, cells were cultured to at 37°C with 5% CO2 until reaching approximately 80% confluency (minimum 48 hrs) and then an aliquot of media was tested according to the manufacturers protocol. Genomic DNA was also extracted (Monarch DNA Isolation Kit, #T3010L) and used for short tandem repeat (STR) authentication, which was performed *via* capillary sequencing (Applied Biosystems). Fingerprints were aligned to reference genomes using the CLASTR algorithm *via* Cellosaurus (https://www.cellosaurus.org/str-search/). All cell lines were expanded for a minimum of five passages prior to metabolite/lipid extraction and LC-MS analysis.

#### Cell seeding for -omics preparation: genomic DNA, RNA, lipids and metabolites

To ensure reproducibility and minimize non-biological variation, cell culture consumables (e.g., FBS, basal media, plates) were derived from the same lot, and their usage was standardized. Cells were seeded at densities ranging from 4.0×10^5^ to 1.5×10^6^ in 10 mL of media in 100 mm dishes and cultured for 48 hrs. Differences in seeding densities accounted for cell size and growth rate to ensure confluency was ∼ 80% after 48 hrs. Each cell line was seeded in replicates of six: one replicate for RNA and gDNA isolations, and 4 to 5 remaining replicates for metabolite and lipid extraction. Cells seeded on 100 mm dishes were harvested 48 hrs post-seeding. COLO205 is a mixed adherent and suspension cell line, so the adherent (COLO205A) and suspension (COLO205S) components were harvested separately and represented as two separate cell lines in the HMA. For adherent cell lines, spent culture media was removed and dishes were rinsed with 7 mL of D-PBS. D-PBS was aspirated and individual 100 mm plates were flash frozen in liquid nitrogen. Frozen 100 mm plates were stored at -80°C until sample preparation. For suspension cell lines, cells were collected in 15 mL Falcon tubes and centrifuged at 300 *rcf* for 5 min. Following centrifugation, spent culture media was removed, cell pellets were washed with 7 mL of DPBS, and subsequently centrifuged again at 300 *rcf* for 5 min. D-PBS was removed from cell pellets after centrifugation, which were then were flash frozen in liquid nitrogen and stored at -80°C until sample preparation.

### RNA sequencing and transcriptomic analysis

RNA was extracted from frozen cell pellets, which were representative samples from cell lines used for LC-MS metabolomics and lipidomic analysis. Briefly, total RNA was isolated using the Monarch Total RNA Miniprep kit (Monarch, #T2010S) with a minimum of 1.0x10^5^ cells and on-column gDNA removal. Quality was assessed *via* bioanalyzer analysis, and a minimum RNA integrity number (RIN) of 7 was used as a cutoff for library preparation and subsequent next-generation sequencing. Libraries were sequenced at a depth of 30 million reads per sample on Novaseq X 25B flow cell (Illumina), paired end 2 x 150bp. Raw reads were mapped to the reference *H. sapiens* genome (GRCh38. p14) and counts were normalized by transcripts per million (TPM).

### Genomic DNA extraction and quantification

Cell pellets for gDNA extraction were concurrently collected with the samples used for metabolomic profiling. Extraction of gDNA was done using the DNeasy Blood and Tissue Kit (Qiagen, Germany) according to manufacturer’s protocols. DNA concentrations were measured using a NanoDrop ONE spectrophotometer (Thermo Fisher Scientific, USA) and Qubit 1X dsDNA BR Assay (Invitrogen, USA).

### Sample preparation for metabolomics and lipidomics analysis

The sample preparation method used for the development of the HMA was adapted from Chan *et al.* (2023) (*21*). Sample extraction was completed in different batches over the course of 5 days in a block-randomized order such that a single replicate from each of the 70 cell lines was extracted on each day of sample preparation. Harvest solution was prepared using H_2_O/methanol (MeOH) (30:225, v/v) spiked with UltimateSPLASH ONE^TM^ (Avanti Polar Lipids, USA) and TruQuant Yeast Extract (IROA Technologies, USA). UltimateSPLASH ONE^TM^ was spiked at 1 µL per 255 µL of harvest solution while 1/36 of a vial of TruQuant Yeast Extract was reconstituted per 255 µL of harvest solution. Harvest solution was added to adherent cell lines frozen on plates or suspension cell lines frozen in 15 mL tubes. The volumes of harvest solution used were normalized to DNA levels derived from an equivalent replicate from each cell line (*60*). Following the addition of harvest solution, adherent cell lines were scraped and 255 µL from each replicate was taken for metabolite and lipid extraction. For suspension cell lines, 255 µL from each replicate was also taken for metabolite and lipid extraction. Once aliquoted, samples were vortexed for 30 s and sonicated for 5 min in a cold-water bath. Subsequently, 750 µL of cold MTBE was added to each sample, followed by vortexing for 30s, and shaking for 10 min at 650 rpm at 4°C on a Thermomixer R (Eppendorf, Germany). Next, 188 µL of cold LC-MS H_2_O was added to each sample, briefly vortexed and centrifuged for 20 min at 16,000 × *g* at 4°C. 700 µL of the upper layer (lipidome) and 250 µL (metabolome) were separated and transferred to new tubes. The extracted lipid/metabolite fractions were subsequently dried down in a vacuum concentrator (Labconco, USA) at 4°C and stored at -80°C until LC-MS analysis.

### LC-MS metabolomics and lipidomics

All metabolome and lipidome profiling experiments were performed on an Orbitrap IQ-X Tribrid mass spectrometer coupled to a Vanquish Horizon UHPLC system (Thermo Fisher Scientific, USA). The metabolome of HMA cell lines were analyzed using two methods. The lipidome of HMA cell lines were analyzed under a single LC-MS method. Samples of a given method were analyzed continuously under a single sequence. Injection order was randomized prior to any LC-MS analysis. Pooled quality control (QC) samples were analyzed every 10 injections to ensure reproducibility. Pooled QCs consisted of an aliquot derived from each cell line analyzed and were made at the start of each LC-MS method. Periodic injections of a random selection of samples were also performed throughout the run to monitor batch effects.

#### Metabolome profiling

In the first method, metabolite extracts were resuspended in 50 µL of 70% acetonitrile (ACN) in H2O (v/v). Samples were subsequently diluted with 50 µL of 80% acetonitrile in 1 mM of medronic acid (MA) (Agilent, Part Number: 5191-4506). This method analyzed samples under negative H-ESI mode with the following source settings: Spray Voltage, -2.5 kV; Sheath Gas, 35 Arb units; Aux gas, 7 Arb units; Sweep Gas, 0 Arb units; Ion Transfer Tube Temp, 300°C; Vaporizer Temp, 275°C. The MS1 acquisition parameters were as follows: Resolution, 120,000; Scan Range, 67-900 *m/z*; RF Lens, 60%; AGC Target, Custom; Maximum Injection Time, 246 ms. The parameters for data dependent MS/MS acquisition were as follows: Data Dependent Mode, Cycle Time; Time between Master Scans, 0.6 s; Isolation Window, 1 *m/z*; Activation Type, HCD; HCD Collision Energy Type, Normalized; HCD Collision Energies (%), 20, 30, 35, 40, 50; Resolution, 30,000. One subsequent MS/MS scan using assisted CID at 15%, 30%, and 45% collision energies was also performed. Mobile phase A consisted of 10 mM ammonium carbonate and 5 µM MA in H_2_O (pH 9). Mobile phase B consisted of 100% ACN. Chromatographic separation was performed using a SeQuant ZIC-pHILIC column (150 mm × 2.1 mm i.d.; 5 µm polymer; MilliporeSigma, USA) attached to a SeQuant ZIC-pHILIC guard column (20 mm × 2.1 mm; MilliporeSigma, USA) with a 2 µL injection. The column temperature was maintained at 45°C. Flow rate for the method was set at 0.170 mL/min during gradient separation steps. The following LC gradient was used: 0-2 min maintain at 90% B, 2-14 min decrease from 90% B to 22% B, 14-15.8 min maintain at 22% B, 15.8-16.2 min increase to 90% B, 16.2-17.5 min maintain at 90% B, 17.5-18.2 min maintain at 90% B and increase flow rate to 0.250 ml/min, 18.0-20.5 min maintain at 90% B with flow rate at 0.250 mL/min, and 20.5-21 min reduce flow rate back to 0.170 mL/min.

In the second method, metabolite extracts were resuspended in 50 µL of 70% ACN in H2O (v/v) and then diluted 1:1 with 50 µL of 70% ACN in H_2_O (v/v). This method analyzed samples under positive H-ESI mode with the following source settings: Spray Voltage, +3.5 kV; Sheath Gas, 50 Arb units; Aux gas, 10 Arb units; Sweep Gas 1, Arb units; Ion Transfer Tube Temp, 325°C; Vaporizer Temp, 350°C. The MS1 acquisition parameters were as follows: Resolution, 120,000; Scan Range, 67-900 *m/z*; RF Lens, 60%; AGC Target, Custom; Maximum Injection Time, 246 ms. MS1 and data-dependent MS/MS acquisition parameters were identical to the first metabolome profiling method. Mobile phase A consisted of 0.1% formic acid in H_2_O. Mobile phase B consisted of 0.1% formic acid in ACN. Chromatographic separation was performed using a Waters ACQUITY UPLC BEH Amide column (100 × 2.1 mm i.d.; 1.7 µm; Waters Corporation, USA). Flow rate was set at 0.400 mL/min while column temperature was maintained at 40°C. The following LC gradient was used: 0-1 min maintain at 99% B, 1-13 min decrease from 99% B to 65% B, 13-16 min decrease from 65% B to 40% B, 16-17 min maintain at 40% B, 17-18 min increase from 40% B to 99% B, and 18-22 min re-equilibrate at 99% B.

#### Lipidome profiling

Lipid extracts were resuspended in 100 µL of ACN:2-propanol (IPA) (50:50; v/v). The method analyzed extracts using a polarity-switching H-ESI method with the following source settings: Spray Voltage, +3.5 kV, -2.5kV; Sheath Gas, 50 Arb units; Aux Gas, 10 Arb units; Sweep gas, 1 Arb unit; Ion Transfer Tube Temp, 300°C; Vaporizer Temp, 350°C. The MS1 acquisition parameters were as follows: Resolution, 60,000; Scan Range, 250-1500 *m/z*; RF Lens, 40%; AGC Target, Standard; Maximum Injection Time Mode, Auto. The data-dependent MS/MS acquisition parameters were as follows: Data Dependent Mode, Cycle Time; Time between Master Scans, 0.4 s; Activation Type, HCD; Collision Energy Mode, stepped; HCD Collision Energy Type, Normalized; HCD Collision Energies (%), 20, 30, 35; Resolution, 15,000. Mobile phase A consisted of ACN:H_2_O (60:40; v/v) with 10 mM ammonium acetate. Mobile phase B consisted of IPA:ACN (90:10; v/v) with 10 mM ammonium acetate. Chromatographic separation was performed using a Waters ACQUITY UPLC CSH C18 column (100 mm × 2.1 mm i.d.; 1.7 µm; Waters Corporation, USA). Flow rate was set at 0.250 ml/min with a column temperature of 65°C. The follow LC gradient was used: 0-1 min maintain at 15% B, 1-3 min increase from 15% B to 30% B, 3-3.5 min increase from 30% B to 48% B, 3.5-12 min increase from 48% B to 82% B, 12- 17 min increase from 82% B to 99% B, 17-18 min maintain at 99% B, 18-18.1 min decrease from 99% B to 15% B, 18.1-22 min maintain at 99% B.

#### MS/MS acquisition using AcquireX

To increase the number of MS/MS spectra, pooled QCs from each LC-MS method were analyzed using the Deep Scan AcquireX workflow in Xcalibur (Thermo Fisher Scientific, USA). At least three repeated injections of the pooled QCs were performed with each LC-MS method. AcquireX parameters were as follows for all three AcquireX runs: Exclusion Override Factor, 3; Exclusion List Peak Window Extension, 0 s; Inclusion List Peak Window Exclusion, 0 s; and Inclusion List Peak Fragmentation Threshold, 50%. Any metabolites still without MS/MS were acquired using a manually curated targeted inclusion list.

### Metabolite identification and peak curation

All metabolites (detected by HILIC-MS methods) were identified at MSI level 1 (*22*, *23*). For MSI level 1 identifications, MS/MS spectra and RT from samples were matched to authentic reference standards. For most lipid classes, lipid mixes were used as reference standards for identification. Lipid standards were also synthesized for lipid classes that did not have reference standards readily available. Remaining lipid species were identified at a minimum confidence level of MSI level 2 as authentic reference standards were not available for every lipid species in the HMA. Accurate mass and MS/MS spectra were used for lipid identification **(Fig. S2)**. MS/MS spectra were matched to predicted MS/MS spectra derived from lipid class information, fatty acid chain length, and double bond content. RT trends were also used to evaluate whether a putative lipid species belonged to a particular lipid class. RT of putative lipid species were predicted based on lipid class information, fatty acid chain length, and double bond content and matched to measured RT. Identified lipids were reported in sum composition notation. Peaks of identified metabolites and lipids were manually integrated using Skyline-daily (v23.1.1.353). Adducts for metabolites and lipids were selected based on peak intensity, peak quality, and group CV. The same adduct was selected for every lipid species in the same lipid class. Peak areas from Skyline were exported and used for downstream data processing.

### Synthesis of lipid standards

UltimateSPLASH ONE lipidomics standards were purchase from Avanti Polar Lipids. Lauric acid (C12:0), myristic acid (C14:0), palmitic acid (C16:0), stearic acid (C18:0), lignoceric acid (C24:0) 3-ketosphinganine, and phytosphingosine were purchased from Cayman Chemicals. Batyl alcohol, diethyl pyrocarbonate (DEPC), dimethylformamide (DMF), dichloromethane (DCM), N,N-dicyclohexylcarbodiimide, and 4-dimethylaminopyridine were purchased from Sigma Aldrich. All reagents used were LC-MS grade purity.

A modified protocol was used for the synthesis of ceramides and their derivatives (*61*). Briefly, 10 mM stock solutions of various fatty acids (C12:0, C14:0, C16:0, C18:0, C24:0) and sphingoid backbones (3-ketosphinganine and phytosphingosine) were prepared in 1:3 DMF:DCM. The following was added to a 900 μL aliquot of 1:3 DMF:DCM equipped with gentle heating and magnetic stirring: 1.75 μL DEPC (11.5 μmol), 1.046 μL of the desired fatty acid stock solutions (10.46 μmol), 1.046 μL of the desired sphingoid backbone (10.46 μmol), and 1.60 μL triethylamine (11.5 μmol), topped to 1 mL total volume. Reaction progress was monitored using thin-layer chromatography with an 18:2 chloroform:MeOH solvent mixture. After 60 minutes, the mixture was evaporated under reduced pressure using a CentriVap and resuspended in 1:1 ACN:IPA.

A modified protocol was used for the synthesis of ether-linked glycerolipids (*62*). Briefly 10 mM stock solutions of various fatty acids (C12, C14, C16, C18, C24) were prepared in chloroform. The following was added to 10.4 mg of batyl alcohol (0.03 mmol) resuspended in 300 μL chloroform equipped with gentle heating and magnetic stirring: 37.14 mg N,N-dicyclohexylcarbodiimide (0.18 mmol) dissolved in 300 μL chloroform, and 3.7 mg 4-dimethylaminopyridine (0.03 mmol) dissolved in 300 μL chloroform, 12 μL of the desired fatty acid stock solutions (0.12 mmol), topped to a 1 mL total volume. Reaction progress was monitored using thin-layer chromatography with an 18:2 chloroform:MeOH solvent mixture. After 60 min, the mixture was evaporated under reduced pressure using a CentriVap and resuspended in 1:1 ACN:IPA. All synthesized lipid standards were validated by using MS/MS fragmentation and RT trends under the same LC-MS method used for lipidomic profiling.

### Derivatization of plasmalogen standards

A modified protocol was used for the mercury-catalyzed cleavage of phospholipids to generate plasmalogen standards (*63*). Briefly, stock solutions of 1% mercury (II) chloride (w/v), 1% sodium chloride (w/v), and 0.1N hydrochloric acid were prepared in water. Lipid standards, including PC (16:1/18:1), PC (P-18:0/18:1), and various phospholipid extracts (Avanti Polar Lipids), were prepared in 1:1 chloroform:methanol. For each reaction, 20 μL of the desired lipid(s) was combined with 1 μL of 1% HgCl_2_ and mixed for 30 s, followed by addition of 204 μL 0.1N HCl. The mixture was stirred at 25°C for 5 min, after which lipids were extracted using an MTBE extraction for LC-MS analysis. Derivatized lipid standards were validated by using MS/MS fragmentation and RT trends under the same LC-MS method used for lipidomic profiling.

### Data handling and normalization

From the exported peak areas, data points that were considered below the limit of detection (LOD) were removed. The LOD was determined for each metabolite and lipid by calculating the mean peak area of blank injections plus three times the standard deviation. Outlier peak areas within a cell line were removed based on an iterative Grubbs’ test. Metabolites and lipid species with >50% of cell line replicates below LOD were considered below LOD. Peak areas for individual metabolites and lipids were normalized to corresponding isotopically labeled internal standards (TruQuant Yeast Extract and UltimateSPLASH ONE^TM^). Metabolites were normalized to the isotopically labeled metabolites found in TruQuant Yeast Extract. Labeled internal standards were used to normalize metabolites when one was available. Surrogate internal standards were used to normalize metabolites that did not have a matching isotopologue. For lipids, peak areas were normalized to a labeled internal standard of the same lipid class from UltimateSPLASH ONE^TM^. All lipid species within the same class were normalized to a corresponding lipid internal standard of the same lipid class. Lipids without a matching lipid class were normalized to a structurally similar surrogate lipid class (e.g., PC O- normalized to PC). Normalized peak areas were calculated by dividing the total peak area of the ^12^C metabolite/lipid by the ^13^C internal standard on a per sample basis. This value was then scaled to the maximum ^13^C peak area for that internal standard across all samples. Peak areas that were below LOD were replaced with the minimum peak area across all samples divided by 2 on a per metabolite/lipid basis. For plots that require a complete dataset (i.e., PCA), remaining missing data points were replaced with values derived from k-nearest neighbors imputation.

### Lectin staining and flow cytometry

Cell lines were cultured as described previously in triplicate. 200,000 cells were seeded into a T25 flask for 48 h prior to harvest. For adherent cell lines, spent culture media was aspirated, and samples were washed with 1 mL of D-PBS. Next, samples were mechanically detached with a scraper and centrifuged at 300 *rcf* for 5 min. Samples were subsequently resuspended in 3 mL of D-PBS until lectin staining. For suspension cell lines, samples were centrifuged at 300 *rcf* for 5 min. Next, samples were subsequently washed in D-PBS and centrifuged again at 300 *rcf* for 5 min. Lastly, samples were resuspended in 3 mL of D-PBS until lectin staining. Cell lines were washed twice with PBS and stained with Zombie NIR viability dye to identify live cells (1:1500, BioLegend #423106). Cells were then washed once with lectin staining buffer (HBSS with Ca^2+^ and Mg^2+^, 1% BSA) in preparation for lectin staining. Cells were stained with FITC conjugated Sambucus Nigra Lectin (SNA) (Invitrogen #L32479) at a dilution of 1:2000. Cells were incubated for 20 minutes on ice followed by a final wash with lectin staining buffer and analysis by flow cytometry. Flow cytometry was performed on a Cytek Aurora (3 lasers, violet (405 nm), blue (488 nm), and red (640 nm)) and acquired data was analyzed using FlowJo software (V10.10.0, BD Biosciences). To account for differences in cell size, FITC median fluorescence intensity (MFI) was normalized to forward scatter area (FSC-A) MFI. Figures were created on GraphPad Prism 10.

### Dose-dependent (1S,3R)-RSL3 cell viability assays

Cell lines were cultured as previously mentioned. 10,000 cells were seeded into a 96-well plate for 24 h. Cells were subsequently treated with a dilution series of (1S,3R)-RSL3 (Cayman Chemical, USA) for 24 h. (1S,3R)-RSL3 dosage started at 10 µM and decreased 10-fold at each dilution. Following treatment, 20 µL of PrestoBlue^TM^ (Thermo Fisher Scientific, USA) was added to each sample. Samples were subsequently incubated at 37°C for 2 h. Fluorescence intensity was measured on a Varioskan LUX microplate reader (Thermo Fisher Scientific, USA). Cell viability was determined as a percentage of RFU from DMSO vehicle control samples. Cell viability curves were created on GraphPad Prism 10.

### Statistical and computational analysis

#### Fatty acid composition heatmaps (FACHs)

FACHs were generated for each lipid class/sample group pairing in the HMA as briefly described in Chan *et al.* (2023) (*21*). For a given lipid class, peak areas of lipid species comprising the class were converted to proportional contribution values by normalizing peak areas to sum to 1 within each sample. Mean proportional contribution values were then computed for each species by averaging proportional contributions across replicate samples. Counts of fatty acyl (FA) carbon chain atoms and double bonds were extracted from each lipid species’ annotation, producing a bi-variate discrete distribution of proportional contribution values. This bi-variate distribution was then plotted as a heatmap, using the number of FA carbon chain length as the x-axis and the number of FA double bonds as the y-axis. Marginal distributions were computed by aggregating and summing proportional contribution values for species sharing either the same FA carbon chain length or FA double bond count and were plotted as marginal bar plots sharing the FACH plot axes. The code for generating FACH plots, implemented in Python, is available at https://github.com/montenegroburkelab/fach. Users can customize output in various ways. For example, users can specify whether to annotate plots with marginal mean FA carbon chain length or double bond counts. These annotations can be added graphically using dashed lines and/or explicitly using text. Users are also given the option to export the normalized data values visualized in the FACHs, allowing for further downstream analysis. To facilitate the comparison of FA compositions between FACHs, color bar scales can be set on a per-sample-group basis, such that proportional contribution values map to the same colors across FACHs of different sample groups.

#### Shannon entropy calculations

An entropy value was calculated for each metabolite in the HMA, allowing for the identification of tissue-specific metabolite enrichment. For a given metabolite, a profile of mean peak areas was computed across cell lines derived from the same tissue type. This profile of mean peak areas was then normalized to sum to 1 and passed as a probability distribution to the *entropy* function from the Python SciPy library to compute an entropy value, *H_m_*, for the tissue distribution of the metabolite *m* (*64*). Here, the natural logarithm was used such that 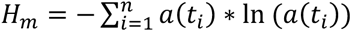 where *a*(*t*_*i*_) represents the mean peak area for metabolite *m* in tissue *t_i_*

#### Statistical analysis

Statistical analyses were conducted in R and Python. Mean metabolite values of corresponding cell lines were used for tissue-based comparisons or correlation analyses. Results from any statistical test were corrected accordingly for multiple testing hypothesis (e.g., Mann Whitney-U test with Benjamini-Hochberg correction).

## Acknowledgements

We would like to acknowledge the Centre for Computational Medicine at the Hospital for Sick Children (Toronto, Canada) for their contributions to the Human Metabolome Atlas online web portal. We would like to thank Dr. Daniel Schramek for the PDC005 cell line.

## Author contributions

Conceptualization: JKC, ATQ, JRMB; Methodology: JKC, NSL, OT, WDG, BYL, VA, ATQ, JRMB; Investigation: JKC, NSL, OT, WDG, BYL, VA, VC, SMA, MM, AJD, ATQ; Visualization: JKC, NSL, BYL, VA, ATQ, JRMB; Formal Analysis: JKC, NSL, OT, BYL, JRMB; Supervision: LJE, JRMB; Funding Acquisition: LJE, JRMB; Writing – original draft: JKC, ATQ, JRMB; Writing – review & editing: JKC, NSL, OT, WDG, BYL, VA, VC, SMA, LJE, ATQ, JRMB.

## Data, code, and materials availability

Materials are available upon request. All data in this manuscript are available in the main text or the supplementary materials. There is also a dedicated resource that allows for extensive visualization of the data described in this manuscript (<u>hma.ccbr.utoronto.ca</u>).

