## Supplementary material for "An atlas of the human metabolome": Suppl. Figs.

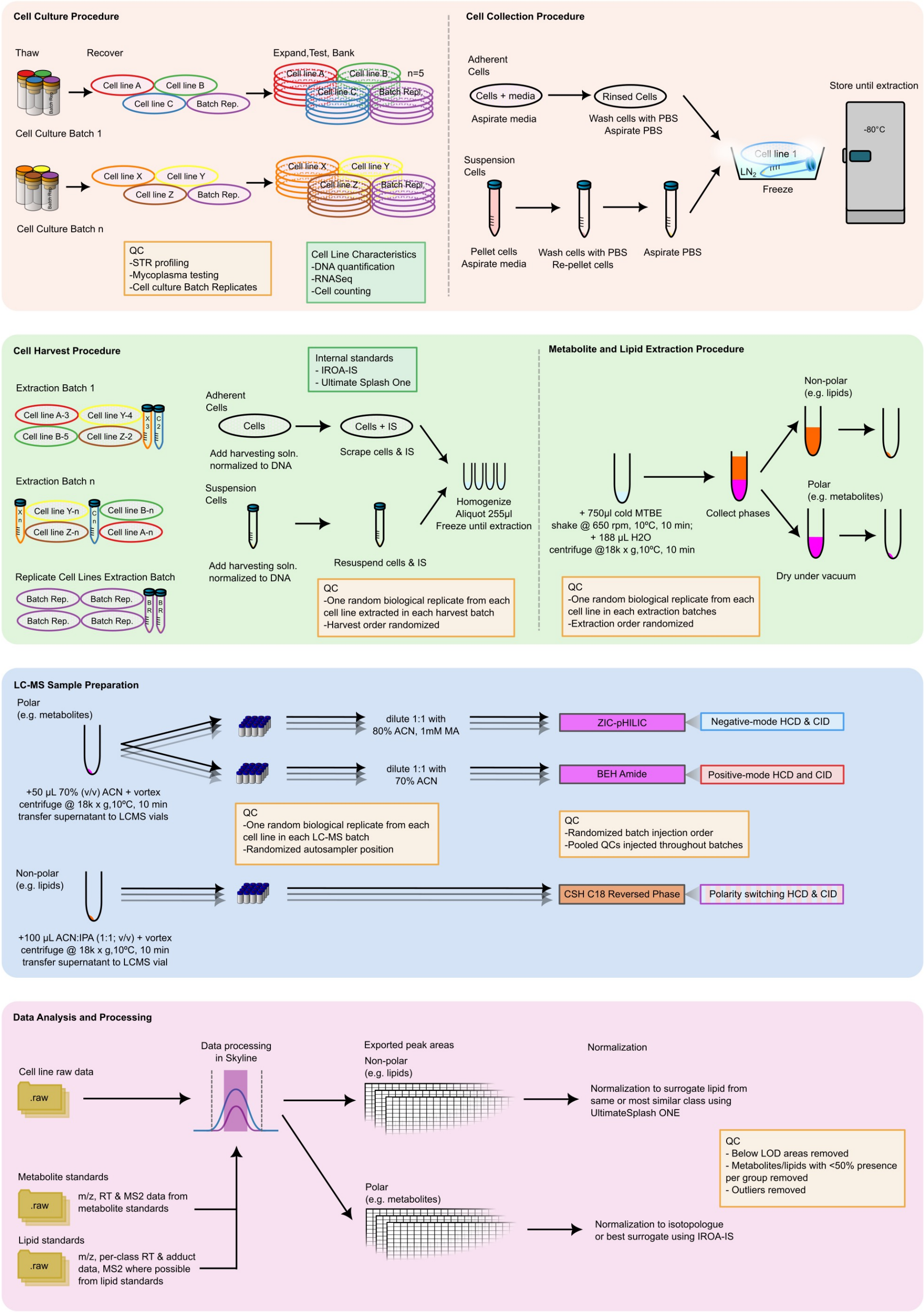


**Fig. S1 Experimental pipeline for the Human Metabolome Atlas.** Prior to cell culture, quality control of cell lines was assured via short-tandem repeat (STR) profiling and mycoplasma testing. From thaw, cell lines were cultured in replicates of up to 5 for 48 hrs. From each replicate, cell lines were washed with PBS and flash frozen in liquid nitrogen. Adherent cell lines were frozen on the original culture dishes while suspension cell lines were pelleted and frozen in tubes. All samples were stored at -80˚C until extraction. Replicates from the HAP1 and U937 cell lines were cultured throughout this stage to control for technical variation in cell culture. For metabolite and lipid extraction, harvest solution (H2O:MeOH, 30:225, v/v) spiked with internal standards were added directly to samples. Volume of harvest solution added per sample was scaled to DNA content. Samples were scraped and/or resuspended in harvest solution and aliquoted. To minimize non-biological variation, individual cell line replicates were distributed across harvest batches in a randomized manner. An adapted biphasic MTBE extraction method was employed to extract metabolites and lipids simultaneously. Following extraction, samples were dried under vacuum and stored at -80˚C until analysis. For LC-MS analysis, three complementary LC-MS methods were employed to maximize metabolite and lipid coverage. Samples were reconstituted in appropriate injection solvents. For analytical quality control, sample injections were randomly distributed across LC-MS batches. Pooled quality control samples and batch control samples were used to monitor instrument stability. For data analysis, raw MS files were manually processed using Skyline. Peak areas were exported and used for downstream data handling and normalization.

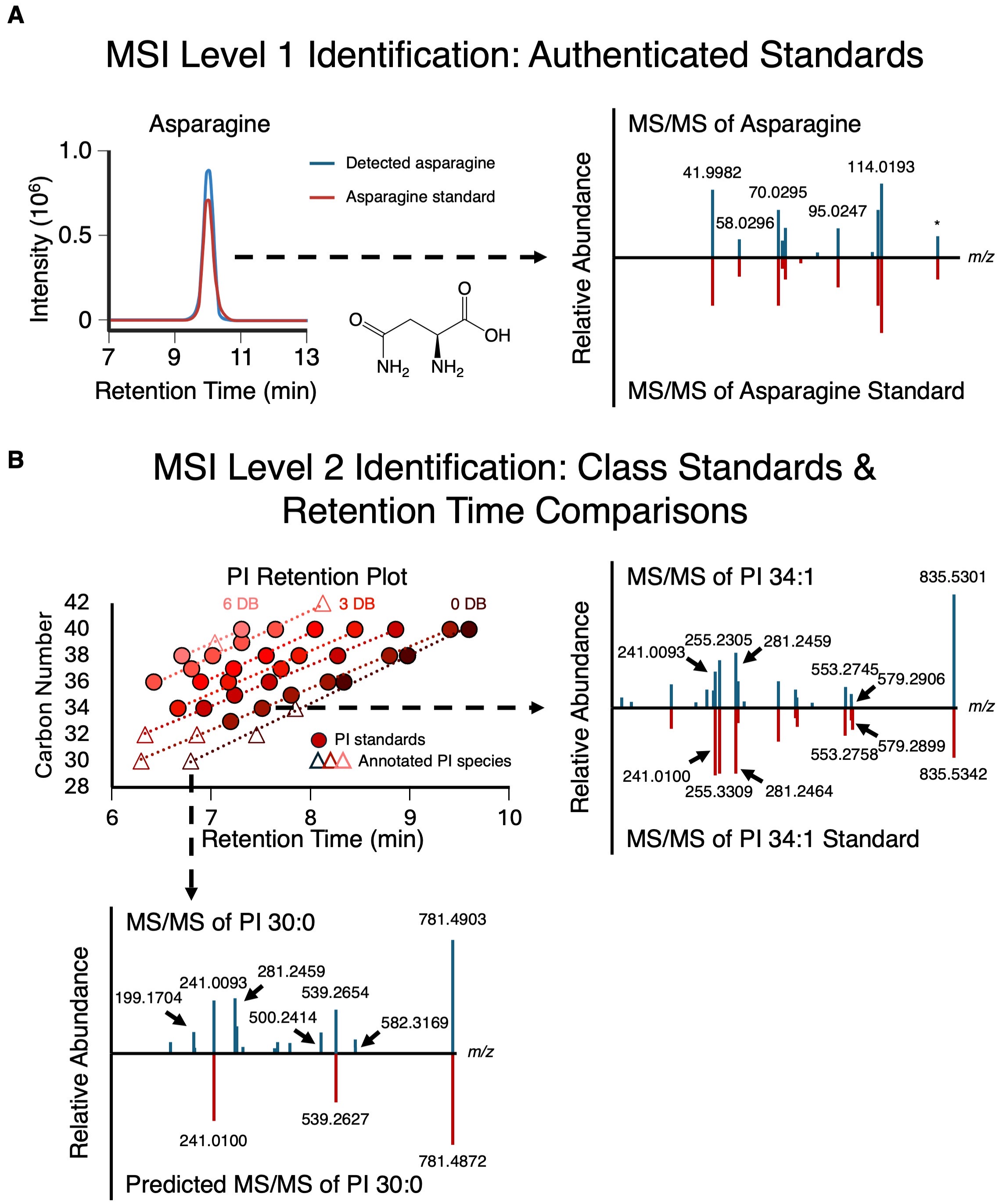


**Fig. S2. Identification of metabolites and lipids in the HMA. (A)** Identification of metabolites at MSI level 1. All polar metabolites were identified at MSI level 1 confidence with matching MS/MS spectra and retention time between measured metabolic feature and authentic standards. **(B)** Identification of lipids at MSI level 2. Lipid species were identified based on accurate mass and MS/MS fragmentation. Retention time (RT) trends were used to validate the identities of lipid species as species of the same class and double bond content tend to elute in a linear manner based on acyl chain length. MS/MS spectra of annotated lipid species were matched to predicted MS/MS spectra derived from known diagnostic fragmentation patterns of various lipid classes.

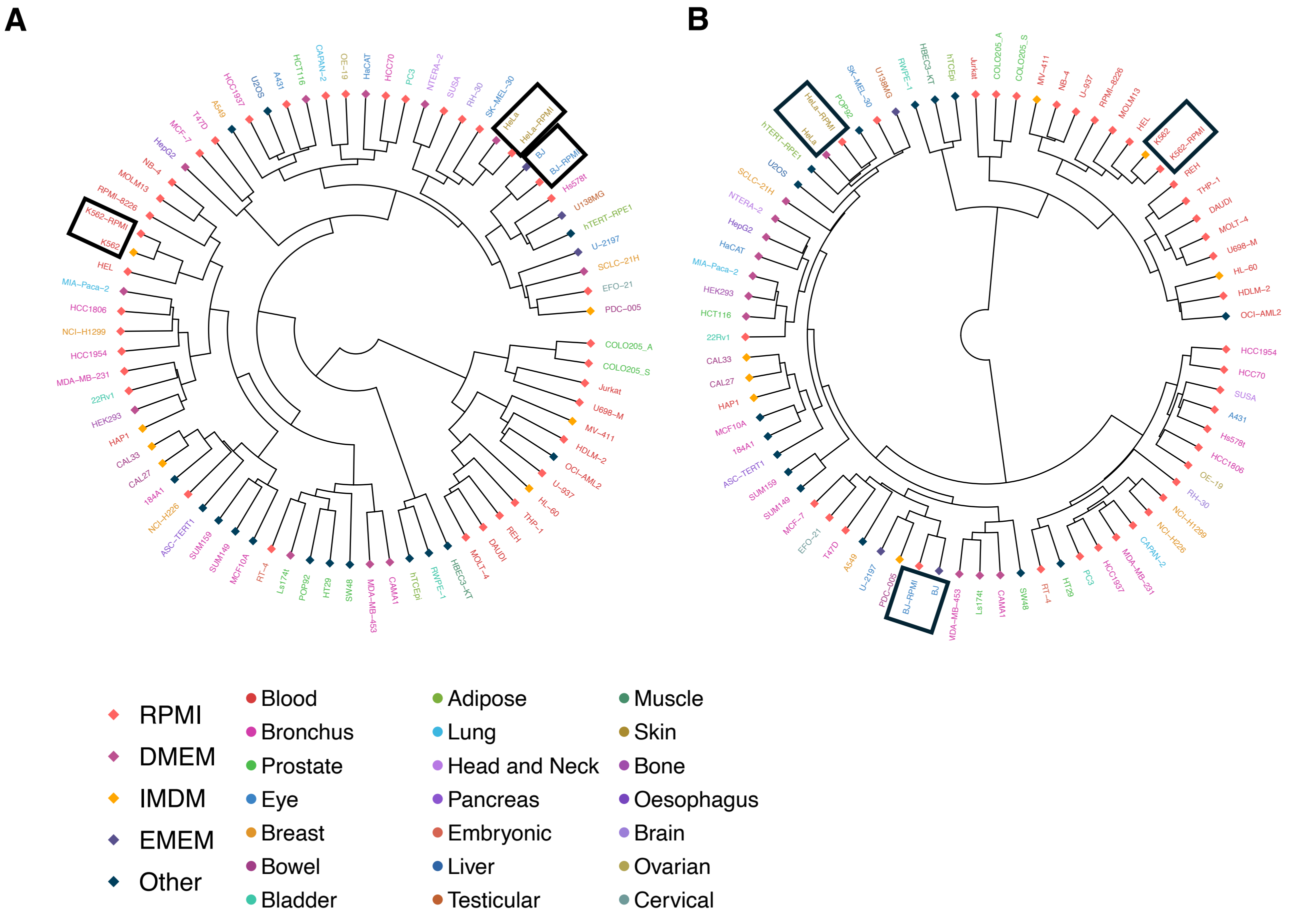


**Fig. S3 Hierarchical clustering of HMA cell lines with media control groups. (A)** Clustered dendrogram of cell lines in the HMA. The data includes the relative levels of all 1768 metabolites and lipids profiled in the HMA. **(B)** Clustered dendrogram of cell lines in the HMA. The data represented is restricted to the relative levels of 200 metabolites. Media controls and corresponding cell lines are highlighted in black boxes. Text color corresponds to cell line tissue of origin. Node colors correspond to basal medium used to grow cell lines.

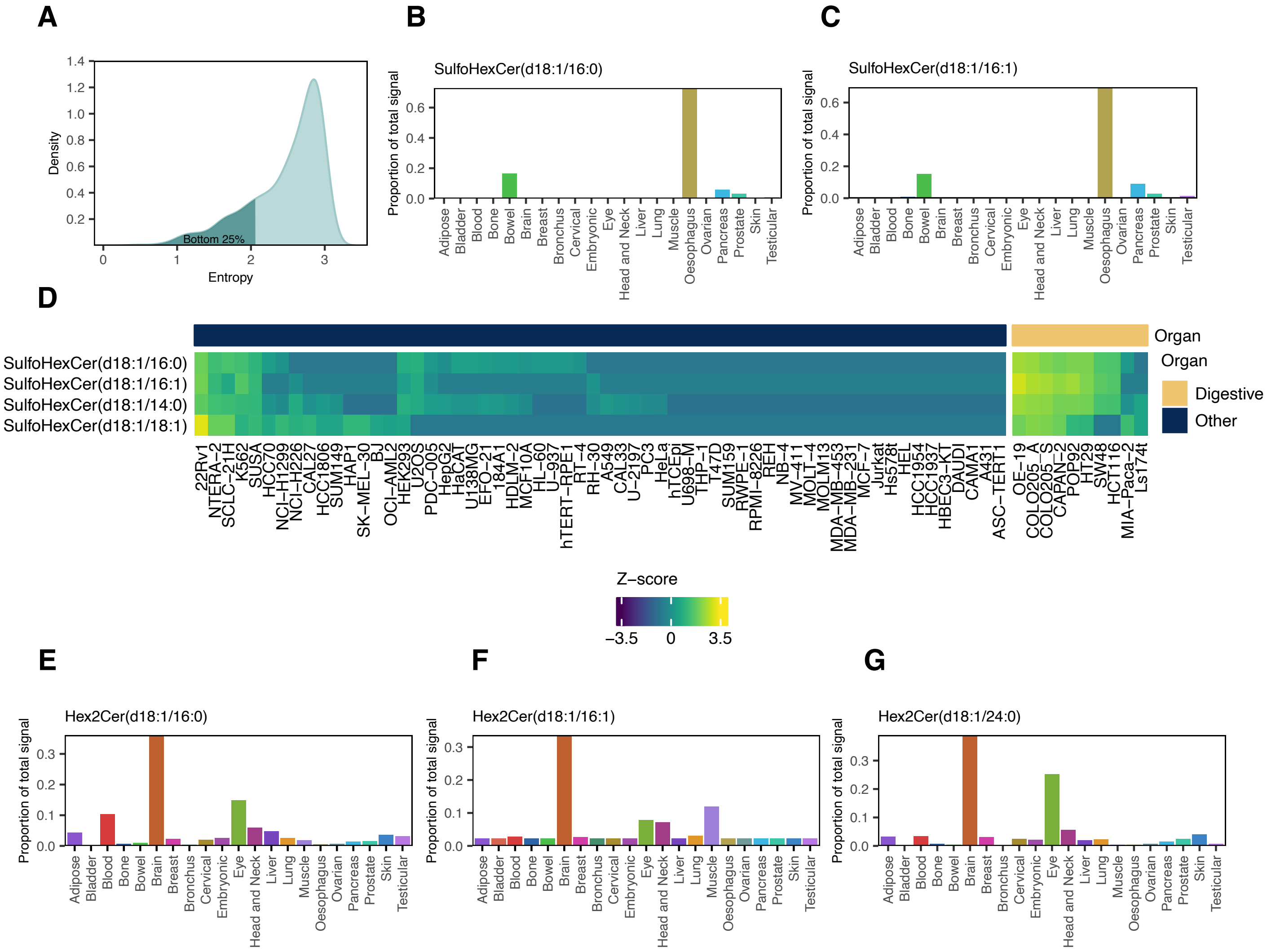


**Fig. S4 Global analysis of lipid profiles in the HMA reveal correlated and tissue-enriched lipid classes. (A)** Density plot highlighting the bottom 25% of metabolites in terms of Shannon's entropy. Lower entropy scores represent more tissue-specific distribution patterns. **(B-C)** Proportional abundance plots for sulfatide (SulfoHexCer) species in the HMA. **(D)** Heatmap of sulfatide species across 70 cell lines. Columns are split based on whether they are related to digestive organs. Data are presented as Z-score of log10-transformed peak areas of sulfatide species. **(E-G)** Proportional abundance plots for dihexosylceramide species (Hex2Cers).

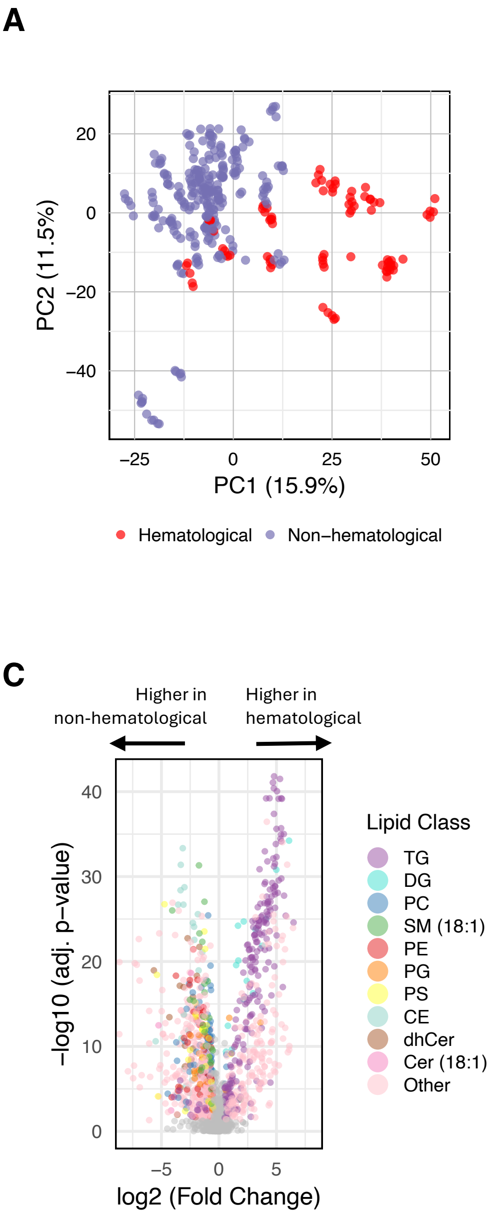


**Fig. S5 Hematological cell lines are characterized by an elevation of nucleotide intermediates and unsaturated TGs. (A)** PCA colored based on hematological or non-hematological tissue of origin. Data presented in the PCA is log-transformed and Z-score scaled. **(B)** Volcano plot depicting differentially abundant metabolites in hematological and non-hematological cell lines. Red points represent metabolites that are elevated in hematological cell lines (log2 fold-change>0.5, adj. p<0.05) while blue points represent metabolites elevated in non-hematological cell lines (log2 fold-change<-0.5, adj. p<0.05). Statistical analyses were performed using Mann-Whitney U test with Benjamini-Hochberg correction. **(C)** Volcano plot depicting differentially abundant lipid species between hematological cell lines and non-hematological cell lines. Significantly different lipids are colored based on the corresponding lipid class. Lipid species that are not significantly different are colored in grey. Significantly different lipids have an absolute log2 fold-change > 0.5 and an adjusted p-value < 0.05 (Mann-Whitney U test, Benjamini-Hochberg correction).

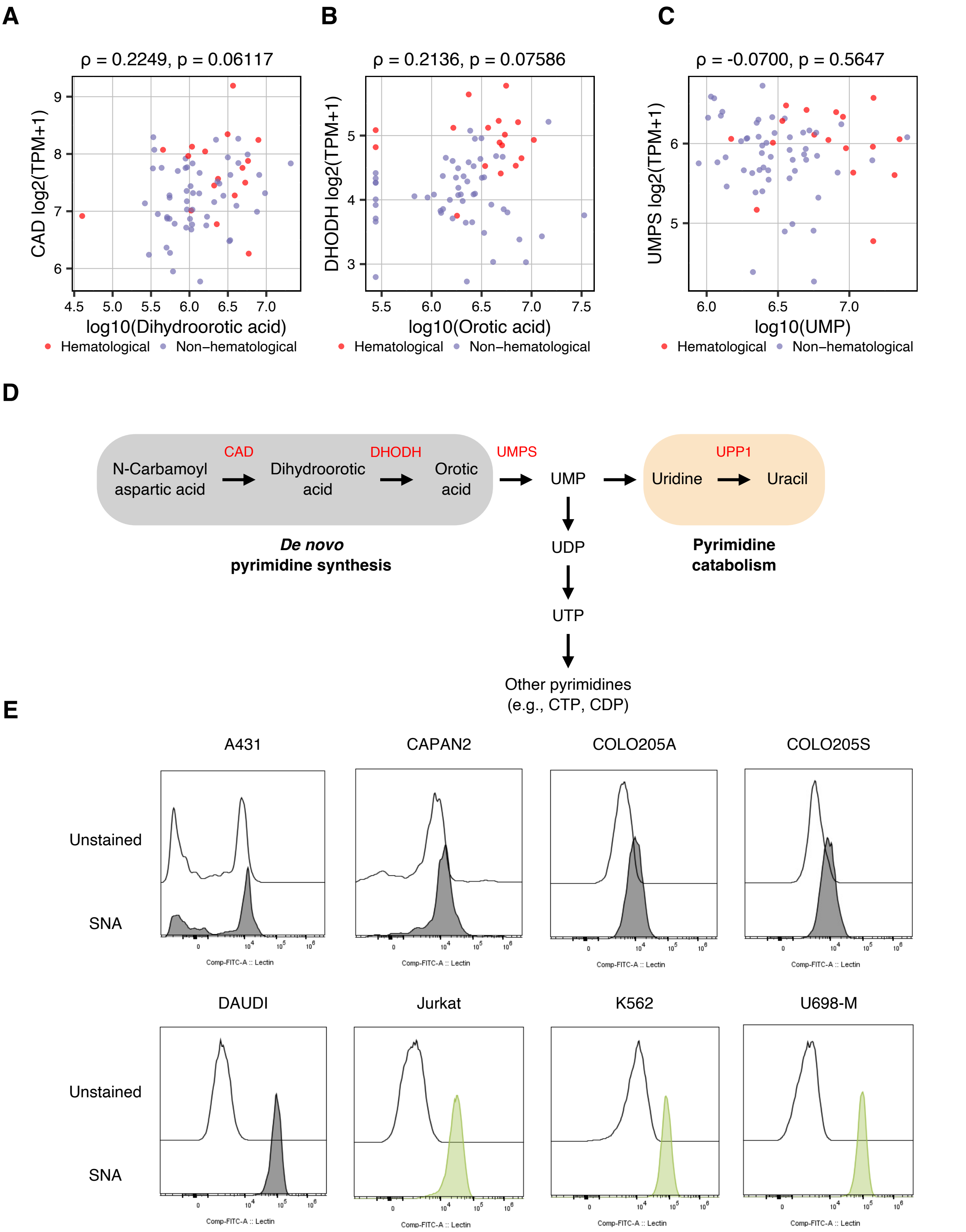


**Fig. S6 Multi-omic integration of HMA metabolomic trends. (A-C)** Scatter plot depicting the Spearman's correlation of pyrimidine intermediates against the expression of corresponding genes of interest. Individual points represent individual cell lines in the HMA. Points are colored based on cell line tissue of origin. **(D)** Schematic for the pathways involved in pyrimidine metabolism. **(E)** Histograms depicting SNA-FITC levels across HMA cell lines.

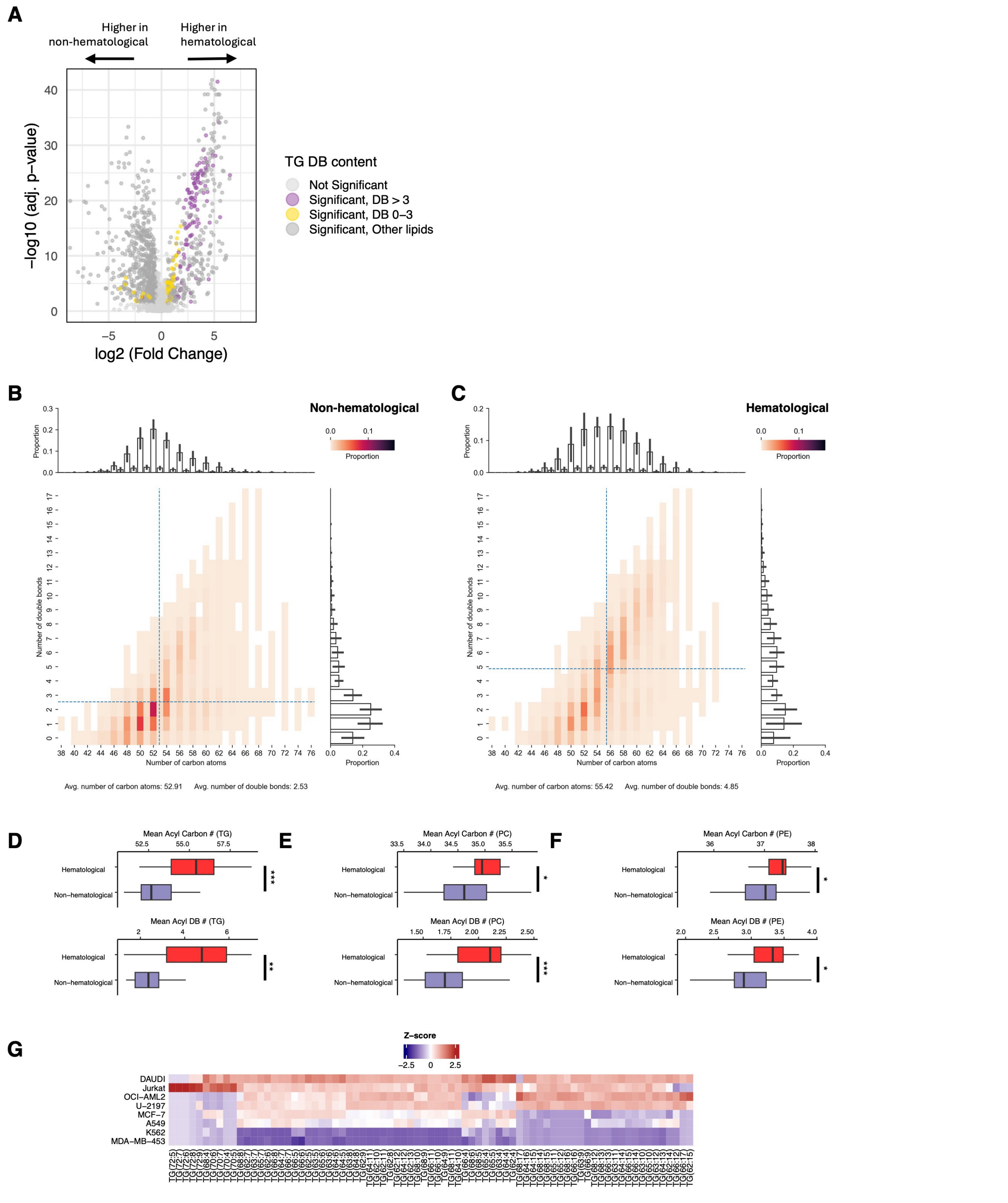


**Fig. S7 Long and unsaturated triglycerides are a lipidomic signature of hematological cell lines. (A)** Volcano plot depicting differentially abundant lipids colored by lipid class and double bond content. Significance is denoted by adjusted p<0.05 (Mann-Whitney U test, Benjamini-Hochberg correction) and absolute log2 fold change>0.5 **(B)** TG FACH of the average non-hematological cell line in the HMA. Data are presented as the proportional abundance of individual TG species. Mean marginals represent the distribution of fatty acyl carbon chain length and fatty acyl double bond number. (**C**) TG FACH plot of the average hematological cell line in the HMA. Data are presented as the proportional abundance of individual TG species. Mean marginals represent the distribution of fatty acyl carbon chain length and fatty acyl double bond number. **(D-F)** Comparison of acyl carbon chain length and acyl double bond number in **(D)** TG, **(E)** PC, and **(F)** PE classes. *, ** or *** denotes significance with p<0.01, p<0.001, and p<0.0001, respectively. Statistical significance was calculated based on Mann Whitney-U test. **(G)** Heatmap of unsaturated TGs. Data are presented as Z-score of log-scaled peak areas.

**A**

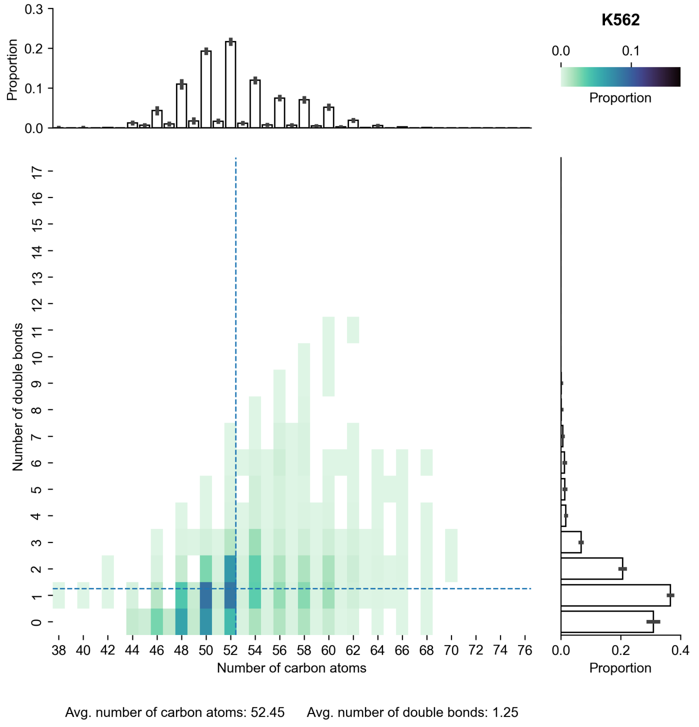


**B**

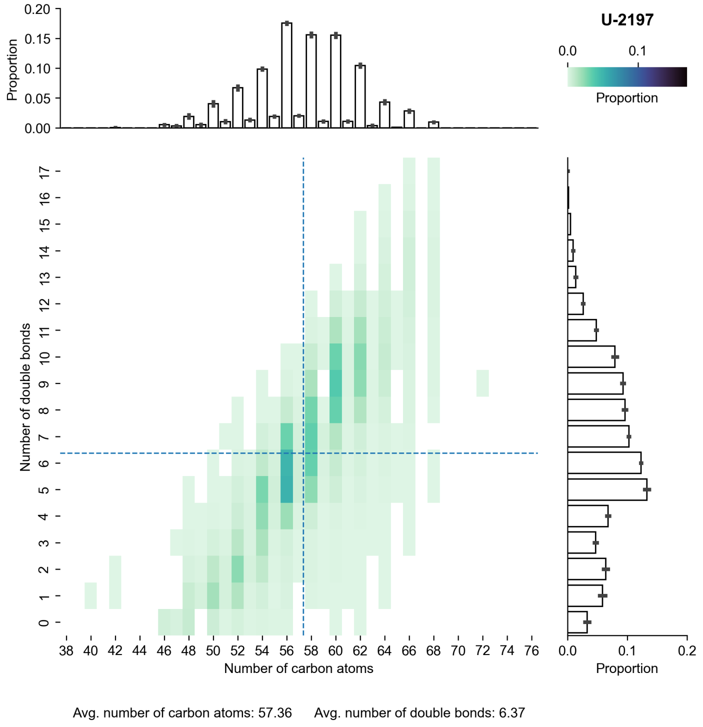


**Fig. S8 TG fatty acid composition of select HMA cell lines.** FACHs of **(A)** K562 and **(B)** U-2197 Data are presented as the proportional abundance of individual TG species. Mean marginals represent the distribution of fatty acyl carbon chain length and fatty acyl double bond number.

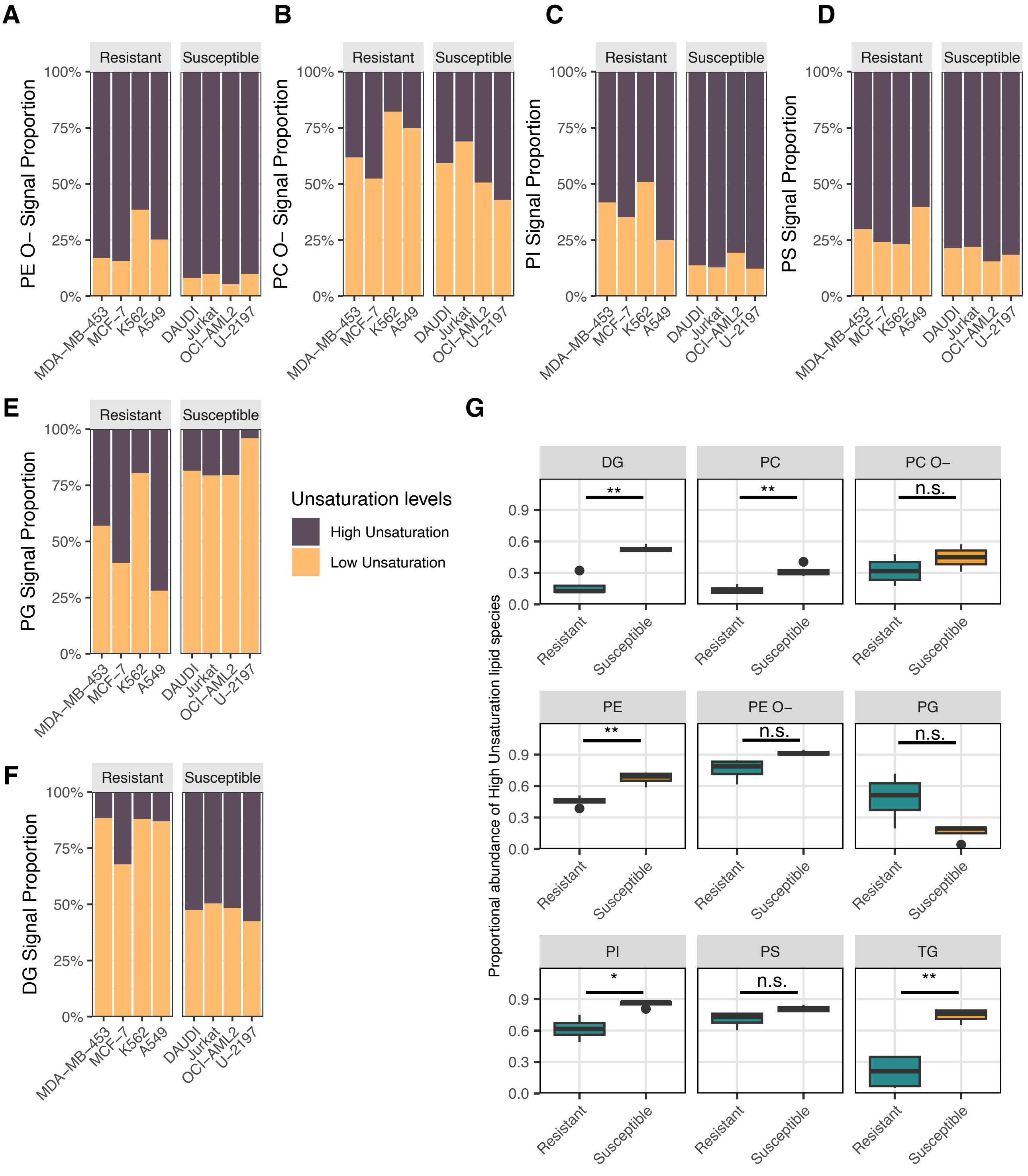


**Fig. S9 Ferroptosis-susceptible cells have a higher proportion of highly unsaturated lipids in specific lipid classes. (A-F)** Stacked bar plots depicting the proportional abundance of (A) PE O-, (B) PC O-, (C) PI, (D) PS, (E) PG, and (F) DG species grouped by unsaturation levels. Abbreviations: Ether-linked phosphatidylethanolamine, PE O-; Ether-linked phosphatidylcholine, PC O-; phosphatidylinositol, PI; phosphatidylserine, PS; phosphatidylglycerol, PG; diglyceride, DG. **(G)** Abundance of highly unsaturated lipid species in RSL3-resistant and susceptible cell lines across glycerolipid and glycerophospholipid classes. Data is presented as proportional abundance of highly unsaturated lipid species for each class. * = p<0.05, ** = p<0.005, n.s. = not significant, by Welch's T test with Benjamini-Hochberg correction.
